# Ketogenic diet synergistic reprogramming of both host and microbiome promotes tissue regeneration

**DOI:** 10.64898/2026.04.11.717958

**Authors:** Motoyoshi Nagai, Victor Band, Liang Chi, Margery Smelkinson, Benjamin Schwarz, Nathan T. Brandes, Andrew Burns, Paula Juliana Perez-Chaparro, Kathryn E. McCauley, Dan Corral, Nicolas Bouladoux, Verena M. Link, Lixin Zheng, Michael G. Constantinides, Michael Otto, Niki M. Moutsopoulos, Yasmine Belkaid

## Abstract

Nutrition influences host physiological processes, yet how diets reshape host physiology, microbial functions, or host–microbe interactions to promote regeneration remains poorly explored. Here, we show that a ketogenic diet (KD), enriched in fats and low in carbohydrates, reprograms both skin microbial and immune functions to promote tissue repair. KD enhances IL-17A activity in γδ T cells and mucosal-associated invariant T (MAIT) cells, accelerating tissue repair, while KD-induced skin lipidomic alterations enhance both the abundance and metabolic output of *Staphylococcus epidermidis*. Metatranscriptomic and lipidomic analyses revealed increased riboflavin biosynthesis and sphingomyelinase (Sph)-dependent ceramide production in *S. epidermidis* under KD conditions. Genetic depletion of microbial *ribD,* a key enzyme for riboflavin biosynthesis, or of *sph* compromised the ability of the bacteria to promote tissue repair. Thus, host nutritional status drives tissue regeneration by synergistically rewiring host and microbial functions, providing new insights into how diet can be harnessed to regulate host physiology.

## Introduction

Diet influences virtually all physiological systems, including those that govern immune cell function (*1*, *2*). Both the quantity and quality of nutrients consumed can directly shape immune responses by altering nutrient availability and immune function. In addition, diet can indirectly affect immunity by modifying the composition and activity of the microbiota, which in turn helps regulate host immunity.

Recent evidence indicates that changes or reductions in dietary intake can modulate the activity of diverse immune cell populations, thereby shaping host immunity in both beneficial and detrimental ways. For example, we and others have shown that dietary restriction enhances T cell function and improves host survival in models of infection (*3*, *4*). This growing understanding of the connection between nutrition and immunity has opened new possibilities for developing dietary interventions that modulate immune responses in diseases such as infection, cancer, and chronic inflammatory disorders (*5*, *6*).

Host nutrition also profoundly affects microbial ecosystems, thereby shaping symbiotic relationships with the host that maintain physiological fitness and immune homeostasis (*7–9*). The microbiota has a remarkable capacity to adapt its composition and function in response to physiological changes, a feature thought to support host resilience (*10–12*). For example, dietary restriction can enhance immune protection by reshaping the gut microbiota and promoting microbiota-derived metabolites that support myeloid cell function (*13*). In addition, studies in both humans and mice have shown that a ketogenic diet (KD), a high-fat, low-carbohydrate diet, can modulate host immune responses through remodeling of the gut microbiota (*14*, *15*).

However, while dietary interventions hold considerable therapeutic promise, how to harness nutrition in a rigorous and mechanistically driven manner remains poorly understood. Moreover, as nutrition can shape both host physiology and microbiota function, how coordinated adaptations of host and microbial activity could converge to promote beneficial outcomes has not been addressed.

The skin represents the largest barrier organ and harbors a specialized immune system (*16*). Skin-resident microbial communities are primarily located within the stratum corneum of the epidermis and play important roles in the development of cutaneous immunity and the maintenance of skin homeostasis (*17–19*). Emerging evidence shows that interactions between the skin microbiota and the immune system are critical regulators of tissue repair following injury or infection (*20–23*). These observations led us to hypothesize that dietary interventions could promote tissue repair by reshaping both microbial and host immune responses in the skin.

Here, we explored whether nutrition could be precisely harnessed to promote one of the most fundamental and vital processes for host survival: tissue repair. Specifically, we investigated whether (1) defined dietary interventions could enhance tissue repair and (2) maximal impact would result from simultaneously reshaping both microbial and host immune functions.

Our results reveal that a defined nutritional intervention promotes tissue repair through coordinated reprogramming of the metaorganism. Notably, we uncover a previously unrecognized role for the ketogenic diet, a widely used dietary approach, in promoting tissue repair. Furthermore, we show that this enhanced tissue repair results from adaptive changes in both host and microbial compartments, which synergistically enhance immune functions and microbial-derived processes associated with tissue regeneration.

## Results

### Ketogenic diet and skin microbiota enhance wound healing

To assess the potential impact of dietary interventions on tissue repair, we altered the nutritional status of mice via dietary restriction or nutrient-modified diets for 4–6 weeks prior to skin biopsy. To assess the potential contribution of the skin microbiota, all mice were also topically associated with *Staphylococcus epidermidis*, a bacterium we have previously shown promotes tissue repair (*22*, *24*, *25*), two weeks before punch biopsy. Tissue repair was quantified 5 days after injury by measuring the length of the epidermal tongue, which reflects proliferating and migrating epithelial fronts (**Fig. 1A**).

**Fig. 1.**
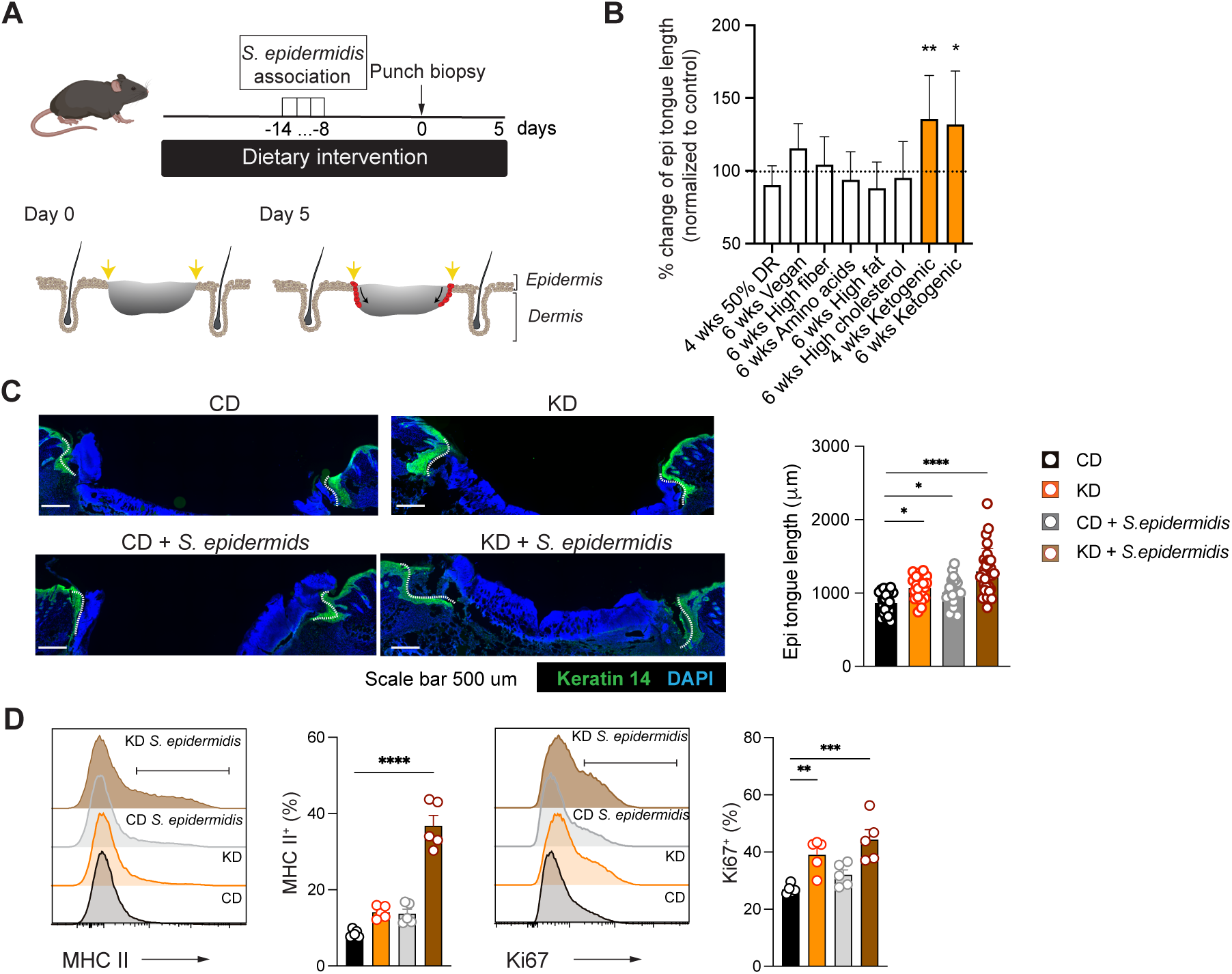
KD and skin microbiota enhance wound healing. (A) Experimental timeline. Mice were topically associated with *S. epidermidis* (LM087) on days −14, −12, −10, and −8, followed by full-thickness punch biopsy on day 0 and analysis on day 5. Dietary status was modified by dietary restriction or nutrient-modified diets. (B) Fold changes in epidermal tongue length in diet-intervention groups relative to *ad libitum*–fed controls or diet-matched control groups. (C–D) Mice were fed a control diet (CD) or ketogenic diet (KD) for 4 weeks and associated with *S. epidermidis* or vehicle (Tryptic Soy Broth, TSB). (C) Representative immunofluorescence images of wounds at day 5. Keratin 14 (green) labels the advancing epidermal tongues (white dashed lines). Scale bars, 500 μm. Right, quantification of epidermal tongue length; each dot represents one epidermal tongue. (D) Histograms of MHC II and Ki67 expression in keratinocytes, with quantification of MHC II⁺ and Ki67⁺ keratinocytes (%). Panel (B–C) shows pooled data from at least two independent experiments; (D) is representative of two independent experiments. (B) One-way ANOVA followed by multiple t-tests; (C–D) one-way ANOVA followed by Dunnett’s test. Data are shown as mean ± s.e.m.; each dot indicates an individual mouse or epidermal tongue. *P < 0.05; **P < 0.01; ***P < 0.001; ****P < 0.0001.

Among the dietary interventions tested, only mice fed a ketogenic diet (KD), a diet enriched in fats and depleted in carbohydrates, showed accelerated healing compared to mice fed a control diet (CD) (**Fig. 1B, C**). Accelerated healing was associated with heightened keratinocyte activation and proliferation, as evidenced by increased MHC II and Ki67 expression in KD + *S. epidermidis* group (**Fig. 1D, and fig. S1**). Of note, while both KD alone and *S. epidermidis* topical association alone accelerated repair compared to CD, the maximal effect was observed when mice were both fed KD and associated with *S. epidermidis,* supporting the idea of a synergistic effect (**Fig. 1C, and fig. S2A**). While the impact of KD + *S. epidermidis* was observed in both males and females, the effect was more pronounced in females (**fig. S2B**).

The observation that the presence of *S. epidermidis* maximized the impact of KD suggests that this impact could result from metabolic and functional rewiring of both host and microbial cells.

### KD feeding enhances the colonization of *S. epidermidis* on the skin

Enhanced microbial impact on tissue repair could result from increased microbial biomass, leading to heightened immune activation and/or metabolic and functional reprogramming of microbial cells.

The skin is a desiccated, nutrient-poor environment (*26*, *27*), and changes in nutrient availability can have major impacts on microbial biomass and function. As such, we tested the possibility that an altered nutrient landscape associated with a lipid-rich diet such as KD could enhance *S. epidermidis* colonization. Notably, S. *epidermidis* burden was increased by approximately 20-fold following KD compared to CD-fed mice (**Fig. 2A**). Increased biomass was not associated with microbial translocation to the regional lymph nodes (**fig. S2C**). *16S rRNA* profiling of the skin microbiota further confirmed that KD-fed mice showed increased colonization, with *S. epidermidis* dominating the microbial landscape (**Fig. 2B**). These findings indicate that KD remodeled the skin environment in a manner that favored *S. epidermidis* colonization.

**Fig. 2.**
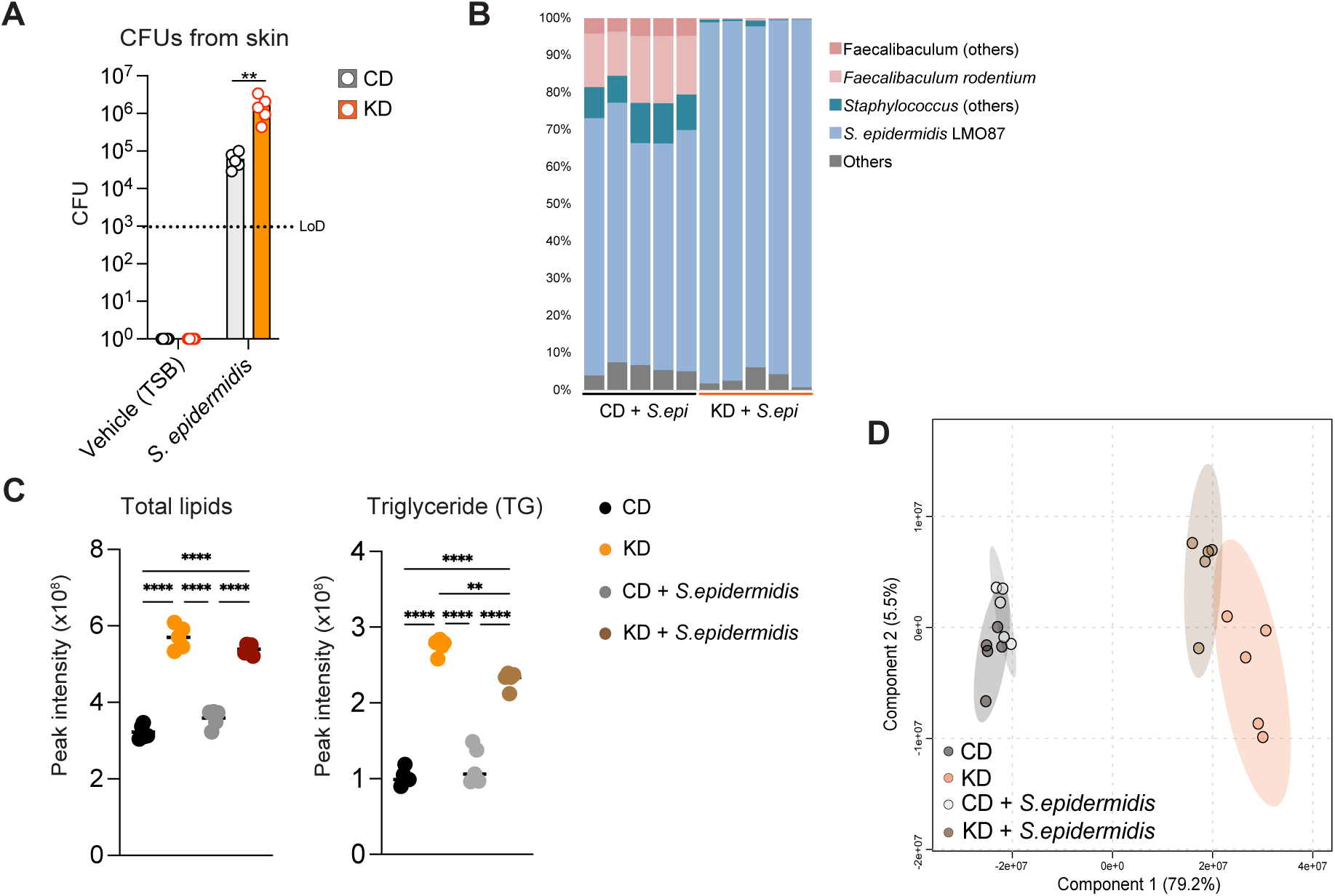
KD feeding enhances the colonization of *S. epidermidis* on the skin. WT mice were fed a CD or KD for four weeks and associated with *S. epidermidis* or vehicle (TSB). (A) Colony-forming units (CFU) of *S. epidermidis* recovered from ear skin. (B) Taxonomic classification of skin microbiota based on 16S rRNA sequencing of back-skin swabs from CD + *S. epidermidis* and KD + *S. epidermidis* mice; each column represents an individual mouse. (C–D) Non-targeted lipidomic analysis of epidermal fractions, including the stratum corneum. (C) Concentrations of indicated lipid species. (D) Principal component analysis (PCA) of epidermal lipid composition; each dot represents an individual mouse. Panel (A) shows representative data from three independent experiments. (A) One-way ANOVA followed by multiple t-tests; (C) one-way ANOVA followed by Tukey’s test. Data are shown as mean ± s.e.m.; each dot indicates an individual mouse. *P < 0.05; **P < 0.01; ***P < 0.001; ****P < 0.0001.

*S. epidermidis* resides within the stratum corneum, a site where microbes can respond to and metabolize host-derived lipids (*17*). To assess whether KD feeding alters the lipid landscape of the outer skin layer, we performed lipidomic profiling of epidermal fractions, which include the stratum corneum, from mice fed CD or KD, with or without *S. epidermidis* colonization. KD feeding markedly increased the total lipid content in the epidermis (**Fig. 2C**), consistent with the high-fat composition of the diet. While diet was the dominant factor shaping epidermal lipid composition, *S. epidermidis* colonization further altered the epidermal lipid milieu (**Fig. 2D**). Of note, the impact of *S. epidermidis* on the epidermal lipid composition was predominantly observed in mice fed KD and not in mice fed CD (**Fig. 2D**).

Triglycerides (TG) constitute a major component of sebum, and *S. epidermidis* can hydrolyze TG into glycerol and free fatty acids through extracellular lipases, using these metabolites as energy sources (*28*, *29*). KD feeding significantly increased epidermal TG abundance, and topical *S. epidermidis* colonization partially reduced TG levels in KD-fed mice (**Fig. 2C**), suggesting active microbial consumption under these lipid-rich conditions.

Thus, alterations in microbial nutrient availability under KD promote microbial colonization and biomass, a phenomenon that can, in turn, enhance local immune activation in a manner that promotes tissue repair.

### KD is associated with metabolic and functional remodeling of pro-repair γδ T cells

To test this possibility, we next assessed host responses to KD. Whole-tissue RNA-seq of the skin showed significant activation of innate and adaptive immune response pathways in the KD + *S. epidermidis* group compared to the CD + *S. epidermidis* group (**Fig. 3A**). This response was not associated with skin inflammation, such as skin thickening or neutrophil infiltration (**fig. S2D, S2E**). Flow cytometric analysis revealed a significant increase in the absolute number of γδ T cells, CD4^+^ T cells, CD8^+^ T cells, ILC3s, and mucosal-associated invariant T (MAIT) cells in the skin of mice fed KD and associated with *S. epidermidis,* compared with CD-fed controls (**Fig. 3B, fig. S1, and fig. S3A**). Particularly, KD + *S. epidermidis* treatment strikingly expanded IL-17A–producing Vγ6⁺ γδ T cells (**Fig. 3B, fig. S3B**).

**Fig. 3.**
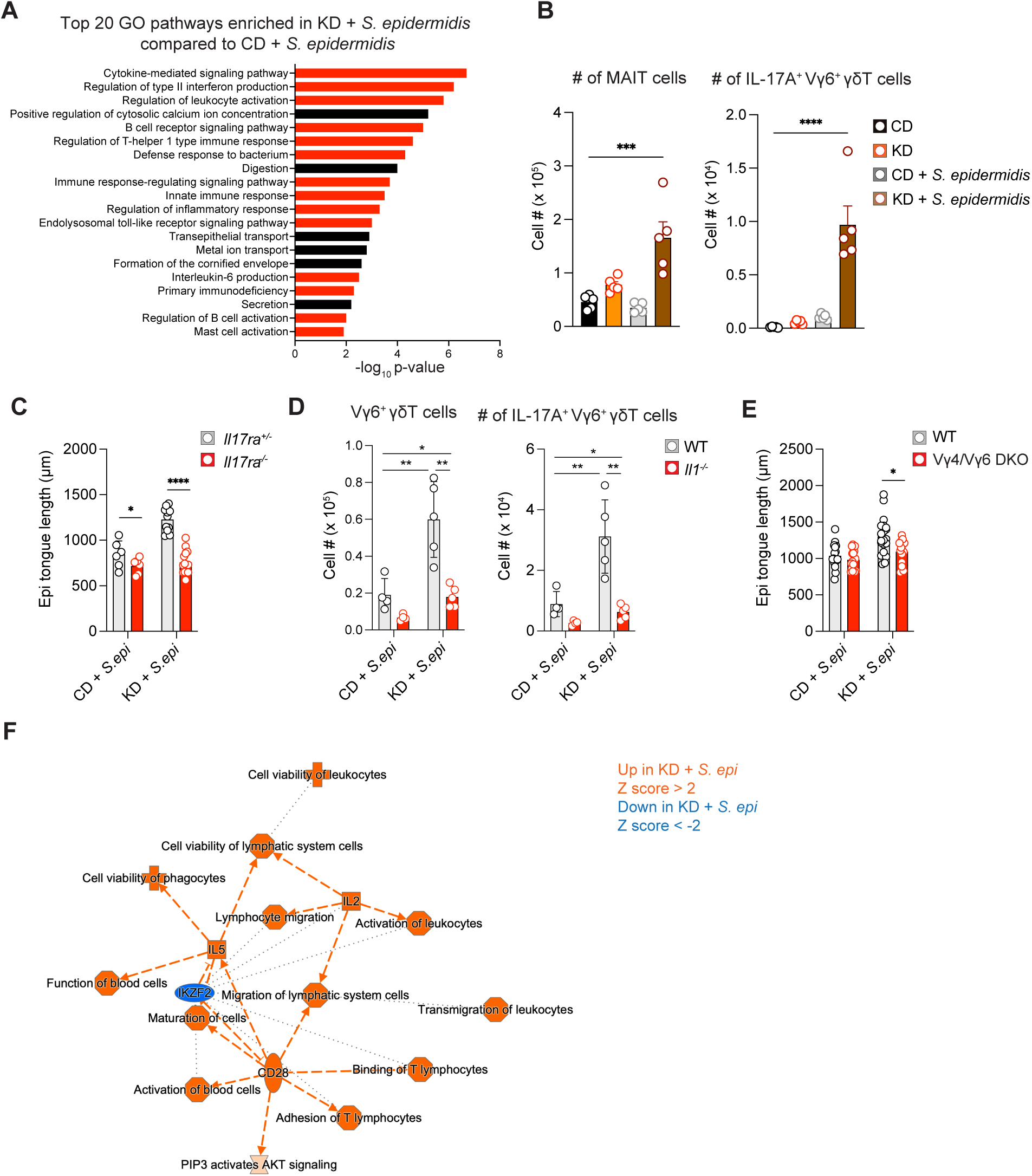
KD is associated with metabolic and functional remodeling of pro-repair γδ T cells. (A,B, F) WT mice were fed a CD or KD for four weeks and associated with *S. epidermidis* or vehicle (TSB). (A) GO enrichment analysis of skin RNA-seq comparing KD + *S. epidermidis* to CD *+ S. epidermidis*; immune-related pathways are highlighted in red. (B) Absolute numbers of indicated immune cell subsets in ear skin. (C, E) *Il17ra^⁺/⁻^* or *Il17ra^⁻/⁻^* mice and Vγ4/Vγ6 double-knockout (DKO) mice or WT mice were fed CD or KD, associated with *S. epidermidis*, and assessed for epidermal tongue length 5 days after wounding. (D) WT and *Il1r^⁻/⁻^* mice were fed CD or KD, associated with *S. epidermidis*, and analyzed for immune cell numbers in ear skin. (F) Bulk RNA-seq of skin Vγ6⁺ γδ T cells from CD + *S. epidermidis* and KD + *S. epidermidis* groups. Ingenuity Pathway Analysis (IPA) identified significantly altered pathways, with upregulated pathways in the KD group shown in orange and downregulated pathways in blue. Panels (B, D) are representative data from at least two independent experiments; panels (C, E) show pooled data from at least two independent experiments. (B) One-way ANOVA followed by Dunnett’s test; (C, E) one-way ANOVA followed by multiple t-tests; (D) one-way ANOVA followed by Tukey’s test. Data are mean ± s.e.m.; each dot represents an individual mouse. *P < 0.05; **P < 0.01; ***P < 0.001; ****P < 0.0001.

We and others have previously shown that IL-17A–producing T cells promote tissue repair through their actions on both the epithelium and the neuronal network (*20*, *23*). In line with this, neither *Il17ra^⁻/⁻^* nor *Il17a^⁻/⁻^* mice exhibited enhanced wound healing following KD and *S. epidermidis* association (**Fig. 3C, and fig. S3C**). Aligned with previous findings that T cell activation in response to *S. epidermidis* was dependent on IL-1α signaling (*19*, *30*), treatment of mice with the IL-1 receptor antagonist anakinra or genetic deficiency of IL-1R abrogated optimal activation of these cells (**Fig. 3D, and fig. S3D**).

Furthermore, consistent with the significant increase in skin γδ T cells, accelerated healing observed in the mice fed KD and associated with *S. epidermidis* was significantly impaired in mice lacking dermal γδ T cell subsets (Vγ4/6 DKO; **Fig. 3E**).

We next performed bulk RNA-seq on sorted skin Vγ6^+^ γδ T cells from CD + *S. epidermidis* and KD + *S. epidermidis* mice (**Fig. 3F**). Ingenuity Pathway Analysis revealed upregulation of TCR signaling pathway (CD28, and IL2) as well as metabolic pathway (PIP3–AKT signaling) in KD + *S. epidermidis* mice compared to CD + *S. epidermidis* (**Fig. 3F**). Using SCENITH, a flow cytometry–based metabolic profiling method (*31*), we found that mitochondrial dependence was significantly increased in Vγ6⁺ γδ T cells from KD + *S. epidermidis* mice compared to CD + *S. epidermidis* group (**fig. S4A, S4B).** In contrast, no differences were observed in glucose dependence (**fig. S4B**). These findings are consistent with previous reports demonstrating that IL-17–producing γδ T cells preferentially rely on oxidative metabolism and exhibit increased mitochondrial activity (*32*). Our data suggest that KD can shape T cell function within the skin thereby broadly amplifying the pro-repair γδ T cell/IL-1/IL-17A axis, a phenomenon that could result from enhanced microbial activity and/or a direct response of T cells to a lipid-rich milieu.

### KD impacts the transcriptional profile of *S. epidermidis*

Bacteria can rapidly sense their surrounding environment and adapt their transcriptional profiles accordingly (*12*, *33*). As such, we tested the possibility that, in addition to enhanced biomass, KD could alter the transcriptional profile of skin-associated bacteria. To this end, we isolated bacterial RNA from the skin of CD + *S. epidermidis* and KD + *S. epidermidis* mice. Consistent with our CFU data, we recovered ∼10-fold more bacterial cDNA from KD + *S. epidermidis* samples (**fig. S5A**). After normalizing the input cDNA, we prepared libraries, performed sequencing, and conducted metatranscriptomic analysis. We identified 8,626 gene families across both groups. Of these, 532 reactions were annotated, and 138 were significantly upregulated in the KD + *S. epidermidis* group compared to the CD + *S. epidermidis* group (**fig. S5B**). Pathway analysis revealed 76 annotated pathways, of which 24 were significantly enriched in KD-associated skin microbiota, including glycolysis, nucleotide biosynthesis, and riboflavin/heme biosynthesis (**Fig. 4A**).

**Fig. 4.**
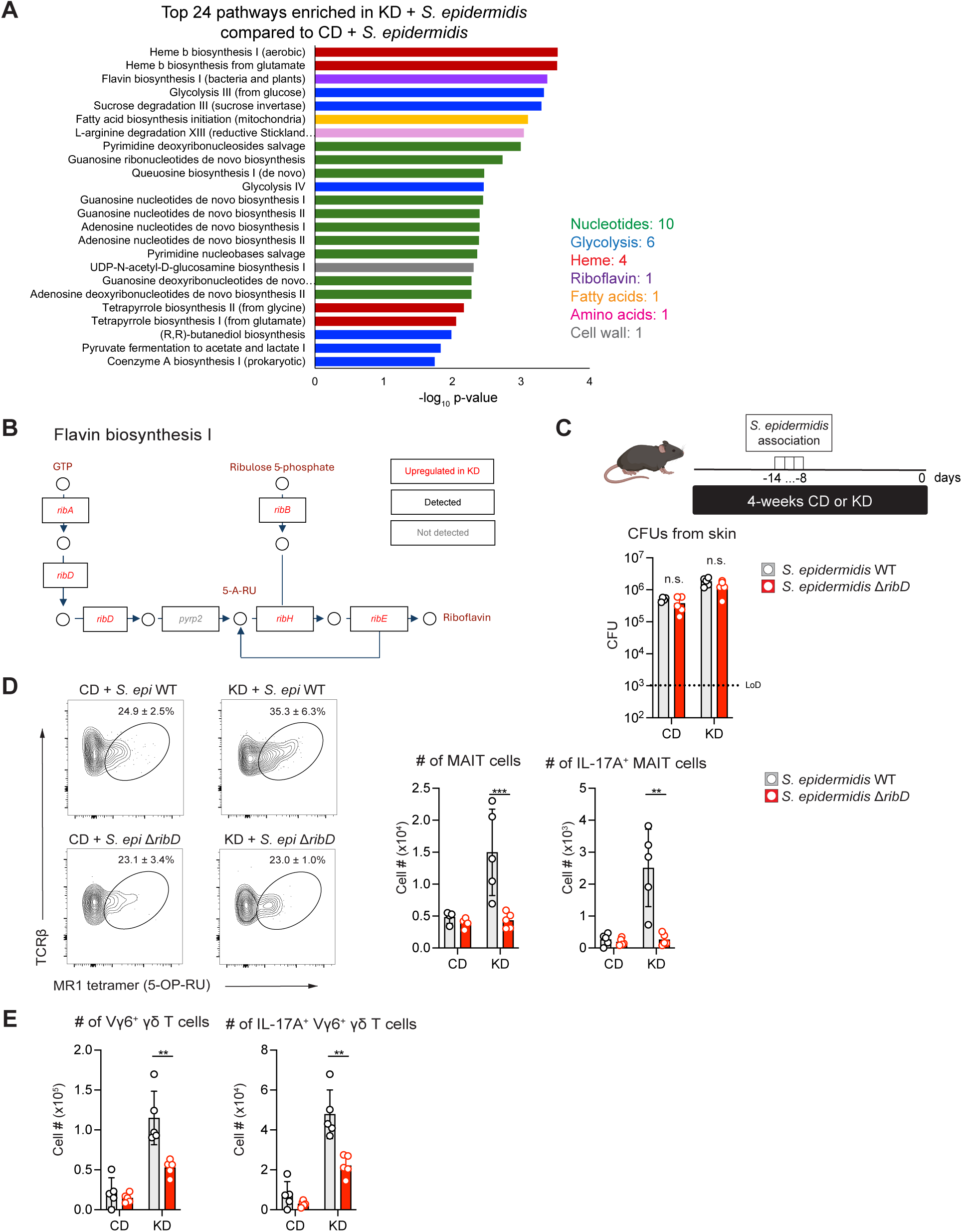
KD impacts the transcriptional profile of *S. epidermidis*. (A) Metatranscriptomic analysis of skin swabs identifying microbial transcriptional changes. 76 annotated pathways were detected, of which 24 were significantly upregulated in KD + *S. epidermidis* mice. The bar graph shows these KD-enriched pathways. (B) Differential expression of flavin biosynthesis genes in skin-associated bacteria from CD *+ S. epidermidis* and KD + *S. epidermidis* mice. Genes significantly upregulated in the KD + *S. epidermidis* group are shown in red. Data are derived from skin metatranscriptomics (related to Fig. 4A and Fig. S5). (C-E) Mice were fed CD or KD and topically associated with *S. epidermidis* LM087 (WT) or the *ribD*-deficient mutant (*ΔribD*). (C) CFU of *S. epidermidis* recovered from skin on day 0. (D) Left, representative contour plots showing MAIT cell gating (live CD45⁺ CD90.2⁺ TCRβ⁺ γδTCR⁻ CD4⁻ CD8β⁻) in ear pinna. Right, absolute number of MAIT cells. (E) Absolute numbers of indicated populations in ear pinna. Abbreviations and EC numbers: *ribA* (EC 3.5.4.25); *ribB* (EC 4.1.99.12); *ribD* (EC 3.5.4.26); *ribE* (EC 2.5.1.9); *ribH* (EC 2.5.1.78); *pyrp2, 5-amino-6-(5-phospho-D-ribitylamino) uracil phosphatase* (EC 3.1.3.104); 5-A-RU, 5-Amino-6-(D-ribitylamino) uracil. Panels (C–E) show representative data from two independent experiments. (C–E) One-way ANOVA followed by multiple t-tests. **P < 0.01; ***P < 0.001.

Glycolysis is a fundamental process by which bacteria generate ATP and increase biomass, thereby supporting their proliferation. Consistent with this, metatranscriptomic analysis detected the complete gene set required for glycolysis (**fig. S5C**). Supporting this, metabolomic profiling showed elevated concentrations of lactate, the end product of glycolysis, in the epidermis of KD + *S. epidermidis* mice (**fig. S5D**), indicating enhanced glycolytic activity under KD conditions.

Several nucleotide biosynthesis pathways, including adenosine deoxyribonucleotide *de novo* biosynthesis, were also significantly upregulated in KD + *S. epidermidis* group (**Fig. 4A, and fig. S6A**). Accordingly, metabolomic profiling of the epidermal fraction revealed increased levels of multiple nucleotides, such as ATP, GTP, and UTP, in KD + *S. epidermidis* mice (**fig. S6B, S6C**). In addition, we observed upregulation of the heme biosynthesis pathway, which may contribute to the production of porphyrins and vitamin B12 (**fig. S6D**). Of interest, previous studies have shown that bacteria-derived extracellular ATP and porphyrins can promote IL-1 production by keratinocytes (*34*, *35*), a phenomenon that may account, at least in part, for the enhanced Vγ6⁺ γδ T cell responses we observed.

### Altered microbial KD-induced metabolism promotes tissue repair

Our previous work showed that *S. epidermidis* can produce riboflavin intermediates, thereby promoting MAIT cell accumulation in the skin and contributing to wound repair (21). Metatranscriptomic analysis revealed that KD promoted significant upregulation of riboflavin biosynthetic pathways in *S. epidermidis* compared to bacteria from mice fed CD (**Fig. 4A, B**). In agreement with this, we observed a significant increase in MAIT cells in KD + *S. epidermidis* mice compared to controls (**Fig. 3B**). RibD is a key enzyme in riboflavin biosynthesis in *S. epidermidis* (**Fig. 4B**) (*21*, *36*). To assess the potential contribution of enhanced riboflavin metabolism to the activation of cutaneous immunity under KD, WT or *ΔribD S. epidermidis* were topically applied to the skin of CD- or KD-fed mice (**Fig. 4C).** *S. epidermidis* skin colonization was increased in KD-fed mice relative to CD-fed mice but was comparable between WT and mutant strains under both conditions (**Fig. 4C**). As expected, MAIT cell numbers were significantly reduced in KD + *ΔribD*–colonized mice compared with KD + WT–colonized mice (**Fig. 4D**). Of note, KD + *ΔribD* mice also displayed reduced numbers of skin Vγ6⁺ γδ T cells (**Fig. 4E**), an observation pointing to the possibility of cross–talk between the two cell subsets or that riboflavin biosynthetic pathway may directly or indirectly control Vγ6⁺ γδ T cells.

To further explore how KD-driven microbial metabolism contributes to tissue repair beyond immune modulation, we examined alterations in epidermal lipid composition. Epidermal lipidomic profiling revealed that ceramides, and more specifically Cer 18:1, were selectively increased in KD + *S. epidermidis* mice (**Fig. 5 A, B**). This phenomenon was specific to KD and not observed in other lipid-rich settings such as those induced by high fat diet (HFD) (**fig. S7A**). Ceramides are lipid molecules that help restore the skin barrier and coordinate repair after injury (*37*, *38*). Recent work has shown that these lipids can be produced not only by the host but also by *S. epidermidis*, which generates ceramides from sphingomyelin through a sphingomyelinase (Sph)-dependent pathway (**Fig. 5C**) (*17*). To test whether bacteria-derived ceramides contribute to KD-induced lipid changes, we employed an *S. epidermidis* Sph-deficient mutant (*Δsph*). Both strains colonized the skin with similar efficiency under CD conditions and enhanced biomass was observed for both strains when mice were fed KD (**Fig. 5D**). In contrast to WT bacteria, *Δsph S. epidermidis* failed to increase epidermal ceramide levels in KD-fed mice (**Fig. 5E, and fig. S7B, S7C**), indicating that elevated ceramides were predominantly result from bacterial Sph activity. We next assessed the potential contribution of bacteria-derived ceramides in tissue repair. WT or *Δsph S. epidermidis* were topically applied two weeks prior to wounding, and epidermal tongue length was measured five days after injury. Under CD conditions, impaired ceramide production by *S. epidermidis* (*Δsph)* significantly affected wound closure, supporting the idea that this bacteria-derived pathway already contributed to tissue repair at baseline (**Fig. 5F**). Further, the enhanced repair observed in KD was also significantly reduced when a mutant strain was applied, compared to the WT strain (**Fig. 5F**). In line with this, *Δsph* colonization was associated with reduced keratinocyte activation (**fig. S7D**). On the other hand, the number of immune cell populations, including Vγ6⁺ γδ T cells, was comparable between WT- and *Δsph*-colonized mice (**fig. S7E, S7F**), supporting the idea that bacterial ceramides promote tissue repair primarily by modulating keratinocyte responses rather than altering skin immune cell composition.

**Fig. 5.**
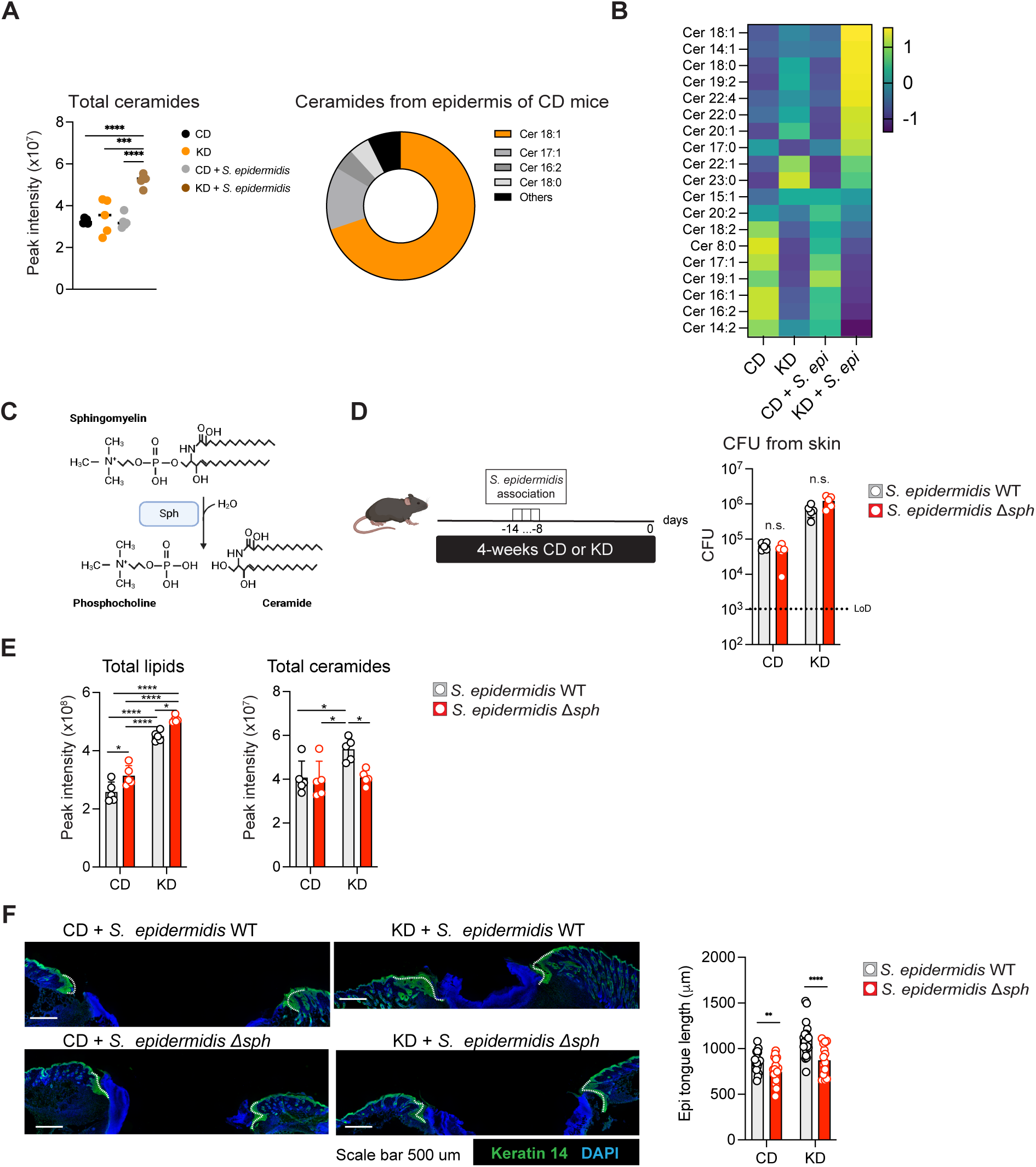
Altered microbial KD-induced metabolism promotes tissue repair. (A, B) Non-targeted lipidomic analysis of epidermal fractions from CD, KD, CD + *S. epidermidis*, and KD + *S. epidermidis* mice. (A) Dot plot showing concentration of indicated lipid. Pie chart showing the relative composition of epidermal ceramide species in the CD group. (B) Heat map showing Z-scored concentrations of various ceramides. (C) Schematic illustrating the activity of *S. epidermidis* sphingomyelinase (Sph), which, together with host sphingomyelinases, converts sphingomyelin into phosphocholine and ceramide. (D-F) WT mice were fed CD or KD and topically associated with S*. epidermidis* 1457 (WT) or the sphingomyelinase-deficient mutant (*Δsph*). (D) CFU of *S. epidermidis* recovered from skin. (E) Levels of total lipids and total ceramides in epidermis measured by lipidomic analysis. (F) Left, representative immunofluorescence images of back-skin wounds 5 days after biopsy. Keratin 14 (green) labels the advancing epidermal tongues; DAPI stains nuclei (blue). Scale bars, 500 μm. Right, epidermal tongue lengths. Each dot represents an individual measurement. Panels (D–E) are representative of two independent experiments; (F) are pooled from two independent experiments. (A, E) One-way ANOVA followed by Tukey’s test; (D, F) one-way ANOVA followed by multiple t-tests. *P < 0.05; **P < 0.01; ***P < 0.001; ****P < 0.0001.

Collectively, these findings reveal that dietary modulation reshapes both skin microbial metabolism and host responses in a synergistic manner, allowing for maximal and coordinated impact on the vital process of regeneration.

## Discussion

Here, we propose that diet can be harnessed to synergistically rewire host and microbiome to promote tissue regeneration. Diet is increasingly recognized as a key determinant of host–microbe interactions, yet its impact on microbial behavior at peripheral barrier sites remains poorly understood. We show that a ketogenic diet (KD) profoundly reshapes the metabolic landscape of the skin, altering both microbial abundance and metabolic output. These alterations modify epithelial and immune responses that collectively enhance tissue repair. By integrating host metabolic status with commensal bacterial metabolism, our findings reveal a diet-sensitive, metabolite-driven axis in which microbial products, including intermediates of the riboflavin pathway and ceramides, act on distinct cutaneous cell types to promote wound healing.

Our results further demonstrate that dietary modulation can selectively amplify cutaneous immune pathways that drive tissue regeneration. KD feeding, in the context *o*f *S. epidermidis* topical association, potentiated the IL-1–IL-17 axis by increasing the abundance and activity of Vγ6⁺ γδ T cells and MAIT cells, both of which have established roles in promoting epithelial proliferation and accelerating tissue repair (*20*, *21*). Although our analyses were limited to bulk RNA-seq and SCENITH, Vγ6⁺ γδ T cells from KD-fed mice displayed modest shifts in immunometabolic features, suggesting a potential contribution of metabolic state to their enhanced responsiveness. Notably, these immune changes occurred without overt inflammation, indicating that dietary inputs can tune pro-repair immunity in the skin without inducing broad inflammatory activation. The selective activation of these immune circuits prompted us to examine whether diet also alters microbial functional activity. We therefore characterized the *in vivo* transcriptional programs of *S. epidermidis* under CD and KD feeding.

Although metatranscriptomics has been increasingly applied to high-biomass sites such as the gut, its use in the skin microbiome field remains limited, largely because the skin harbors relatively few microbes and yields low microbial biomass (*39*). Only a handful of studies have performed metatranscriptomic profiling of skin- or wound-associated communities, including acne lesions, burn wounds, and diabetic foot ulcers, and, more recently, healthy human skin (*40–43*). Here, we extend the still limited application of metatranscriptomics in skin research by providing, to our knowledge, the first *in vivo* transcriptional profiling of a defined skin commensal under different dietary conditions. KD feeding not only increased *S. epidermidis* biomass but also rewired its metabolic activity, enhancing glycolysis, nucleotide biosynthesis, riboflavin-pathway activity, and bacterial ceramide production. These results show that diet can remodel the cutaneous metabolic landscape in ways that directly modify microbial function with relevance to tissue repair. A limitation of our study is that microbial RNA could not be recovered from SPF mice not associated with *S. epidermidis*, preventing analysis of how CD and KD remodel endogenous skin microbiota transcription. Improving RNA recovery from low-biomass skin communities will be essential for future work.

The metabolic pathways upregulated in *S. epidermidis* under KD feeding provide mechanistic clues as to how microbial activity may shape host responses. Enhanced glycolysis and nucleotide biosynthesis in *S. epidermidis* under KD conditions may increase extracellular ATP (*44*), a well-established trigger of keratinocyte IL-1 production (*45*), thereby providing a plausible mechanism linking microbial metabolic activity to the augmented IL-1–IL-17 axis observed in KD-fed mice. Upregulation of the riboflavin biosynthetic pathway suggests increased production of intermediates capable of modulating host immunity. Such intermediates are known to activate MAIT cells through MR1 (*36*) and may also influence γδ T cells by enhancing TCR-dependent signaling. Consistent with this possibility, bulk RNA-seq of Vγ6⁺ γδ T cells showed elevated expression of TCR signaling pathways in KD-fed mice. However, whether the same intermediates that stimulate MAIT cells, such as 5-OP-RU, contribute to γδ T cell activation remains uncertain, and the antigenic ligands and presenting elements that engage Vγ6⁺ γδ T cells *in vivo* have yet to be defined. These gaps underscore the need to clarify the precise microbial metabolites and antigen-recognition pathways through which diet-modulated microbes tune γδ T cell responses.

Our data indicate that *S. epidermidis*–derived ceramides promote wound repair through epithelial-intrinsic pathways. KD feeding markedly increased epidermal ceramides, dominated by Cer 18:1, with smaller increases in saturated species such as C18:0. Because long-chain saturated ceramides can activate the keratinocyte receptor FPR2 (*46*, *47*), this minor saturated fraction may contribute to epithelial activation. Ceramides may also enhance repair by improving epidermal hydration and barrier integrity (*17*, *48*). These mechanisms are consistent with our observations that *Δsph S. epidermidis* impaired wound healing without altering immune populations, and that KD increased ceramides in the absence of overt inflammation. Thus, microbial ceramides likely act directly on keratinocytes to support wound repair, complementing the KD-enhanced immune pathways described above.

Together, our study shows that dietary status can reprogram the metabolic activity of a dominant skin commensal, thereby modifying epithelial and immune pathways that promote tissue repair. By revealing that KD feeding enhances specific microbial metabolites, including glycolytic products, nucleotide precursors, riboflavin-pathway intermediates, and ceramides, we identify diet as a key regulator of cutaneous host–microbe interactions. These findings establish a framework in which microbial metabolism links host nutrition to barrier immunity and tissue regeneration and highlight the potential of dietary or microbiota-directed interventions to modulate skin physiology.

## Acknowledgments

This work was supported in part by the Division of Intramural Research of NIAID. The contributions of the NIH authors are considered Works of the United States Government. The findings and conclusions presented in this paper are those of the author(s) and do not necessarily reflect the views of the NIH or the U.S. Department of Health and Human Services. We thank Kimberly Beacht, Ejae Lewis, Eduard Ansaldo, Jessie Polanco, and the NIAID animal facility for technical support; Rafael Arguello for providing SCENITH kit; the NIH Tetramer Core Facility (NIH Contract 75N93020D00005 and RRID:SCR_026557) for providing MR-1/5-OP-RU and CD1d/α-GalCer Tetramer; Shruti Naik, Eric Van Dang, Pamela L Schwartzberg, Dominic P Golec, Keisuke Nagao, Ana Taijeiro García-Quijada, all the members of the Belkaid and Moutsopoulos Laboratories for helpful discussions and for providing constructive feedback on this project. We used ChatGPT (OpenAI) to assist with English language editing. The authors take full responsibility for the content of the manuscript. Schematic figures were created with BioRender (https://biorender.com).

## Author contributions

Conceptualization: YB, MN

Methodology: YB, MN, VB, LC, PJP-C, DC

Investigation: MN, VB, LC, MS, BS, NTB, AB, PJP-C, NB, LZ

Visualization: MN, MS, KM, VML

Resources: MGC, MO, PJP-C

Funding acquisition: YB, NM

Supervision: YB, MN, NM

Writing – original draft: YB, MN

Writing – review & editing: YB, MN

## Data and materials availability

Mutant *S. epidermidis* (*Δsph*) is available from M. Otto under a material agreement with NIAID. RNA-seq, 16S rRNA-seq, and metatranscriptomics raw data were deposited in NCBI BioProject under accession number PRJNA1442133. All data needed to evaluate the conclusions are available in the main text or the supplementary materials. Other datasets (flow cytometry, lipidomic, and metabolomic dataset) and materials are available upon request. Requests should be sent to corresponding authors, Yasmine Belkaid, or Motoyoshi Nagai

## References

1. N. Collins, Y. Belkaid, Control of immunity via nutritional interventions. Immunity 55, 210–223 (2022).

2. T. Okawa, M. Nagai, K. Hase, Dietary Intervention Impacts Immune Cell Functions and Dynamics by Inducing Metabolic Rewiring. Front. Immunol. 11 (2021).

3. N. Collins, S.-J. Han, M. Enamorado, V. M. Link, B. Huang, E. A. Moseman, R. J. Kishton, J. P. Shannon, D. Dixit, S. R. Schwab, H. D. Hickman, N. P. Restifo, D. B. McGavern, P. L. Schwartzberg, Y. Belkaid, The Bone Marrow Protects and Optimizes Immunological Memory during Dietary Restriction. Cell 178, 1088–1101.e15 (2019).

4. C. Palma, C. La Rocca, V. Gigantino, G. Aquino, G. Piccaro, D. Di Silvestre, F. Brambilla, R. Rossi, F. Bonacina, M. T. Lepore, M. Audano, N. Mitro, G. Botti, S. Bruzzaniti, C. Fusco, C. Procaccini, V. De Rosa, M. Galgani, C. Alviggi, A. Puca, F. Grassi, T. Rezzonico-Jost, G. D. Norata, P. Mauri, M. G. Netea, P. de Candia, G. Matarese, Caloric Restriction Promotes Immunometabolic Reprogramming Leading to Protection from Tuberculosis. Cell Metab 33, 300–318.e12 (2021).

5. C. Wilhelm, J. Surendar, F. Karagiannis, Enemy or ally? Fasting as an essential regulator of immune responses. Trends Immunol 42, 389–400 (2021).

6. A. H. Lee, V. D. Dixit, Dietary Regulation of Immunity. Immunity 53, 510–523 (2020).

7. E. Ansaldo, T. K. Farley, Y. Belkaid, Control of Immunity by the Microbiota. Annu Rev Immunol 39, 449–479 (2021).

8. P. Zhang, Influence of Foods and Nutrition on the Gut Microbiome and Implications for Intestinal Health. Int J Mol Sci 23, 9588 (2022).

9. A. L. Kau, P. P. Ahern, N. W. Griffin, A. L. Goodman, J. I. Gordon, Human nutrition, the gut microbiome and the immune system. Nature 474, 327–336 (2011).

10. T. Okada, S. Fukuda, K. Hase, S. Nishiumi, Y. Izumi, M. Yoshida, T. Hagiwara, R. Kawashima, M. Yamazaki, T. Oshio, T. Otsubo, K. Inagaki-Ohara, K. Kakimoto, K. Higuchi, Y. I. Kawamura, H. Ohno, T. Dohi, Microbiota-derived lactate accelerates colon epithelial cell turnover in starvation-refed mice. Nat Commun 4, 1654 (2013).

11. J. L. Sonnenburg, F. Bäckhed, Diet-microbiota interactions as moderators of human metabolism. Nature 535, 56–64 (2016).

12. T. Ferenci, Sensing nutrient levels in bacteria. Nat Chem Biol 3, 607–608 (2007).

13. S.-J. Han, A. Stacy, D. Corral, V. M. Link, M. K. De Siqueira, L. Chi, A. Teijeiro, D. S. Yong, P. J. Perez-Chaparro, N. Bouladoux, A. I. Lim, M. Enamorado, Y. Belkaid, N. Collins, Microbiota configuration determines nutritional immune optimization. Proceedings of the National Academy of Sciences 120, e2304905120 (2023).

14. V. M. Link, P. Subramanian, F. Cheung, K. L. Han, A. Stacy, L. Chi, B. A. Sellers, G. Koroleva, A. B. Courville, S. Mistry, A. Burns, R. Apps, K. D. Hall, Y. Belkaid, Differential peripheral immune signatures elicited by vegan versus ketogenic diets in humans. Nat Med 30, 560–572 (2024).

15. Q. Y. Ang, M. Alexander, J. C. Newman, Y. Tian, J. Cai, V. Upadhyay, J. A. Turnbaugh, E. Verdin, K. D. Hall, R. L. Leibel, E. Ravussin, M. Rosenbaum, A. D. Patterson, P. J. Turnbaugh, Ketogenic Diets Alter the Gut Microbiome Resulting in Decreased Intestinal Th17 Cells. Cell 181, 1263–1275.e16 (2020).

16. C. Zhang, G. R. Merana, T. Harris-Tryon, T. C. Scharschmidt, Skin immunity: dissecting the complex biology of our body’s outer barrier. Mucosal Immunol 15, 551–561 (2022).

17. Y. Zheng, R. L. Hunt, A. E. Villaruz, E. L. Fisher, R. Liu, Q. Liu, G. Y. C. Cheung, M. Li, M. Otto, Commensal *Staphylococcus epidermidis* contributes to skin barrier homeostasis by generating protective ceramides. Cell Host & Microbe 30, 301–313.e9 (2022).

18. T. Nakatsuji, J. Y. Cheng, R. L. Gallo, Mechanisms for control of skin immune function by the microbiome. Current Opinion in Immunology 72, 324–330 (2021).

19. S. Naik, N. Bouladoux, J. L. Linehan, S.-J. Han, O. J. Harrison, C. Wilhelm, S. Conlan, S. Himmelfarb, A. L. Byrd, C. Deming, M. Quinones, J. M. Brenchley, H. H. Kong, R. Tussiwand, K. M. Murphy, M. Merad, J. A. Segre, Y. Belkaid, Commensal–dendritic-cell interaction specifies a unique protective skin immune signature. Nature 520, 104–108 (2015).

20. P. Konieczny, Y. Xing, I. Sidhu, I. Subudhi, K. P. Mansfield, B. Hsieh, D. E. Biancur, S. B. Larsen, M. Cammer, D. Li, N. X. Landén, C. Loomis, A. Heguy, A. N. Tikhonova, A. Tsirigos, S. Naik, Interleukin-17 governs hypoxic adaptation of injured epithelium. Science 377, eabg9302 (2022).

21. M. G. Constantinides, V. M. Link, S. Tamoutounour, A. C. Wong, P. J. Perez-Chaparro, S.-J. Han, Y. E. Chen, K. Li, S. Farhat, A. Weckel, S. R. Krishnamurthy, I. Vujkovic-Cvijin, J. L. Linehan, N. Bouladoux, E. D. Merrill, S. Roy, D. J. Cua, E. J. Adams, A. Bhandoola, T. C. Scharschmidt, J. Aubé, M. A. Fischbach, Y. Belkaid, MAIT cells are imprinted by the microbiota in early life and promote tissue repair. Science 366, eaax6624 (2019).

22. O. J. Harrison, J. L. Linehan, H.-Y. Shih, N. Bouladoux, S.-J. Han, M. Smelkinson, S. K. Sen, A. L. Byrd, M. Enamorado, C. Yao, S. Tamoutounour, F. Van Laethem, C. Hurabielle, N. Collins, A. Paun, R. Salcedo, J. J. O’Shea, Y. Belkaid, Commensal-specific T cell plasticity promotes rapid tissue adaptation to injury. Science 363, eaat6280 (2019).

23. M. Enamorado, W. Kulalert, S.-J. Han, I. Rao, J. Delaleu, V. M. Link, D. Yong, M. Smelkinson, L. Gil, S. Nakajima, J. L. Linehan, N. Bouladoux, J. Wlaschin, J. Kabat, O. Kamenyeva, L. Deng, I. Gribonika, A. T. Chesler, I. M. Chiu, C. E. L. Pichon, Y. Belkaid, Immunity to the microbiota promotes sensory neuron regeneration. Cell 186, 607–620.e17 (2023).

24. J. L. Linehan, O. J. Harrison, S.-J. Han, A. L. Byrd, I. Vujkovic-Cvijin, A. V. Villarino, S. K. Sen, J. Shaik, M. Smelkinson, S. Tamoutounour, N. Collins, N. Bouladoux, A. Dzutsev, S. P. Rosshart, J. H. Arbuckle, C.-R. Wang, T. M. Kristie, B. Rehermann, G. Trinchieri, J. M. Brenchley, J. J. O’Shea, Y. Belkaid, Non-classical Immunity Controls Microbiota Impact on Skin Immunity and Tissue Repair. Cell 172, 784–796.e18 (2018).

25. D. S. Lima-Junior, S. R. Krishnamurthy, N. Bouladoux, N. Collins, S.-J. Han, E. Y. Chen, M. G. Constantinides, V. M. Link, A. I. Lim, M. Enamorado, C. Cataisson, L. Gil, I. Rao, T. K. Farley, G. Koroleva, J. Attig, S. H. Yuspa, M. A. Fischbach, G. Kassiotis, Y. Belkaid, Endogenous retroviruses promote homeostatic and inflammatory responses to the microbiota. Cell 184, 3794–3811.e19 (2021).

26. E. A. Grice, J. A. Segre, The skin microbiome. Nat Rev Microbiol 9, 244–253 (2011).

27. A. L. Byrd, Y. Belkaid, J. A. Segre, The human skin microbiome. Nat Rev Microbiol 16, 143–155 (2018).

28. S. Edupuganti, L. Parcha, L. Mangamoori, Purification and Characterization of Extracellular Lipase from Staphylococcus epidermidis (MTCC 10656). J App Pharm Sci, 057–063 (2017).

29. G. Pablo, A. Hammons, S. Bradley, J. E. Fulton, Characteristics of the Extracellular Lipases from *Corynebacterium Acnes* and *Staphylococcus Epidermis*. Journal of Investigative Dermatology 63, 231–238 (1974).

30. S. Naik, N. Bouladoux, C. Wilhelm, M. J. Molloy, R. Salcedo, W. Kastenmuller, C. Deming, M. Quinones, L. Koo, S. Conlan, S. Spencer, J. A. Hall, A. Dzutsev, H. Kong, D. J. Campbell, G. Trinchieri, J. A. Segre, Y. Belkaid, Compartmentalized Control of Skin Immunity by Resident Commensals. Science 337, 1115–1119 (2012).

31. R. J. Argüello, A. J. Combes, R. Char, J.-P. Gigan, A. I. Baaziz, E. Bousiquot, V. Camosseto, B. Samad, J. Tsui, P. Yan, S. Boissonneau, D. Figarella-Branger, E. Gatti, E. Tabouret, M. F. Krummel, P. Pierre, SCENITH: A flow cytometry based method to functionally profile energy metabolism with single cell resolution. Cell Metab 32, 1063–1075.e7 (2020).

32. N. Lopes, C. McIntyre, S. Martin, M. Raverdeau, N. Sumaria, A. C. Kohlgruber, G. J. Fiala, L. Agudelo, L. Dyck, H. Kane, A. Douglas, S. Cunningham, H. Prendeville, R. Loftus, C. Carmody, P. Pierre, M. Kellis, M. Brenner, R. J. Argüello, B. Silva-Santos, D. J. Pennington, L. Lynch, Distinct metabolic programs established in the thymus control effector functions of γδ T cell subsets in tumor microenvironments. Nat Immunol 22, 179–192 (2021).

33. K. Y. Le, S. Dastgheyb, T. V. Ho, M. Otto, Molecular determinants of staphylococcal biofilm dispersal and structuring. Front Cell Infect Microbiol 4, 167 (2014).

34. M. Wickersham, S. Wachtel, T. Wong Fok Lung, G. Soong, R. Jacquet, A. Richardson, D. Parker, A. Prince, Metabolic Stress Drives Keratinocyte Defenses against *Staphylococcus aureus* Infection. Cell Reports 18, 2742–2751 (2017).

35. K.-J. Spittaels, K. van Uytfanghe, C. C. Zouboulis, C. Stove, A. Crabbé, T. Coenye, Porphyrins produced by acneic Cutibacterium acnes strains activate the inflammasome by inducing K+ leakage. iScience 24, 102575 (2021).

36. L. Kjer-Nielsen, O. Patel, A. J. Corbett, J. Le Nours, B. Meehan, L. Liu, M. Bhati, Z. Chen, L. Kostenko, R. Reantragoon, N. A. Williamson, A. W. Purcell, N. L. Dudek, M. J. McConville, R. A. J. O’Hair, G. N. Khairallah, D. I. Godfrey, D. P. Fairlie, J. Rossjohn, J. McCluskey, MR1 presents microbial vitamin B metabolites to MAIT cells. Nature 491, 717–723 (2012).

37. W. Huang, J. Liu, L. Zhao, H. He, Function of ceramides in the skin and its relationship with skin disease. The Journal of Steroid Biochemistry and Molecular Biology 254, 106842 (2025).

38. J.-H. Lee, J.-H. Kim, T.-I. Hyeon, K.-T. Min, S.-Y. Lee, H.-C. Ko, H.-S. Choi, K.-Y. Ju, Y.- S. Cho, T.-J. Yoon, C24 Ceramide Lipid Nanoparticles for Skin Wound Healing. Pharmaceutics 17, 242 (2025).

39. T. M. Santiago-Rodriguez, B. Le François, J. M. Macklaim, E. Doukhanine, E. B. Hollister, The Skin Microbiome: Current Techniques, Challenges, and Future Directions. Microorganisms 11, 1222 (2023).

40. T. Ojala, A. Lindford, K. Savijoki, H. Lagus, J. Tommila, A. Medlar, P. Kuusela, P. Varmanen, L. Holm, J. Vuola, E. Kankuri, M. Kankainen, Metatranscriptomic assessment of burn wound infection clearance. Clin Microbiol Infect 27, 144–146 (2021).

41. M. Radzieta, T. J. Peters, H. G. Dickson, A. J. Cowin, L. A. Lavery, S. Schwarzer, T. Roberts, S. O. Jensen, M. Malone, A metatranscriptomic approach to explore longitudinal tissue specimens from non-healing diabetes related foot ulcers. APMIS 130, 383–396 (2022).

42. D. Kang, B. Shi, M. C. Erfe, N. Craft, H. Li, Vitamin B12 modulates the transcriptome of the skin microbiota in acne pathogenesis. Sci Transl Med 7, 293ra103 (2015).

43. M. Chia, A. H. Q. Ng, A. Ravikrishnan, A. N. Mohamed Naim, S. Wearne, J. Common, N. Nagarajan, Skin metatranscriptomics reveals a landscape of variation in microbial activity and gene expression across the human body. Nat Biotechnol, 1–12 (2025).

44. I. Hironaka, T. Iwase, S. Sugimoto, K. Okuda, A. Tajima, K. Yanaga, Y. Mizunoe, Glucose triggers ATP secretion from bacteria in a growth-phase-dependent manner. Appl Environ Microbiol 79, 2328–2335 (2013).

45. Y. Ahn, J. Seo, E. J. Lee, J. Y. Kim, M.-Y. Park, S. Hwang, A. Almurayshid, B. J. Lim, J.- W. Yu, S. H. Oh, ATP-P2X7–Induced Inflammasome Activation Contributes to Melanocyte Death and CD8+ T-Cell Trafficking to the Skin in Vitiligo. Journal of Investigative Dermatology 140, 1794–1804.e4 (2020).

46. H. Lin, C. Ma, K. Cai, L. Guo, X. Wang, L. Lv, C. Zhang, J. Lin, D. Zhang, C. Ye, T. Wang, S. Huang, J. Han, Z. Zhang, J. Gao, M. Zhang, Z. Pu, F. Li, Y. Guo, X. Zhou, C. Qin, F. Yi, X. Yu, W. Kong, C. Jiang, J.-P. Sun, Metabolic signaling of ceramides through the FPR2 receptor inhibits adipocyte thermogenesis. Science 388, eado4188 (2025).

47. M. Lebtig, J. Scheurer, M. Muenkel, J. Becker, E. Bastounis, A. Peschel, D. Kretschmer, Keratinocytes use FPR2 to detect Staphylococcus aureus and initiate antimicrobial skin defense. Front. Immunol. 14 (2023).

48. Y. Uchida, Ceramide signaling in mammalian epidermis. Biochim Biophys Acta 1841, 453–462 (2014).

49. D. Mack, M. Nedelmann, A. Krokotsch, A. Schwarzkopf, J. Heesemann, R. Laufs, Characterization of transposon mutants of biofilm-producing Staphylococcus epidermidis impaired in the accumulative phase of biofilm production: genetic identification of a hexosamine-containing polysaccharide intercellular adhesin. Infect Immun 62, 3244–3253 (1994).

50. S. Conlan, L. A. Mijares, NISC Comparative Sequencing Program, J. Becker, R. W. Blakesley, G. G. Bouffard, S. Brooks, H. Coleman, J. Gupta, N. Gurson, M. Park, B. Schmidt, P. J. Thomas, M. Otto, H. H. Kong, P. R. Murray, J. A. Segre, Staphylococcus epidermidis pan-genome sequence analysis reveals diversity of skin commensal and hospital infection-associated isolates. Genome Biol 13, R64 (2012).

51. B. E. Keyes, S. Liu, A. Asare, S. Naik, J. Levorse, L. Polak, C. P. Lu, M. Nikolova, H. A. Pasolli, E. Fuchs, Impaired Epidermal to Dendritic T Cell Signaling Slows Wound Repair in Aged Skin. Cell 167, 1323–1338.e14 (2016).

52. B. J. Callahan, P. J. McMurdie, M. J. Rosen, A. W. Han, A. J. A. Johnson, S. P. Holmes, DADA2: High-resolution sample inference from Illumina amplicon data. Nat Methods 13, 581–583 (2016).

53. N. Weber, D. Liou, J. Dommer, P. MacMenamin, M. Quiñones, I. Misner, A. J. Oler, J. Wan, L. Kim, M. Coakley McCarthy, S. Ezeji, K. Noble, D. E. Hurt, Nephele: a cloud platform for simplified, standardized and reproducible microbiome data analysis. Bioinformatics 34, 1411–1413 (2018).

54. M. Chuvochina, J. Gerken, M. Frentrup, Y. Sandikci, R. Goldmann, H. M. Freese, M. Göker, J. Sikorski, P. Yarza, C. Quast, J. Peplies, F. O. Glöckner, L. C. Reimer, SILVA in 2026: a global core biodata resource for rRNA within the DSMZ digital diversity. Nucleic Acids Res 54, D334–D341 (2026).

55. F. Beghini, L. J. McIver, A. Blanco-Míguez, L. Dubois, F. Asnicar, S. Maharjan, A. Mailyan, P. Manghi, M. Scholz, A. M. Thomas, M. Valles-Colomer, G. Weingart, Y. Zhang, M. Zolfo, C. Huttenhower, E. A. Franzosa, N. Segata, Integrating taxonomic, functional, and strain-level profiling of diverse microbial communities with bioBakery 3. Elife 10, e65088 (2021).

56. A. Blanco-Míguez, F. Beghini, F. Cumbo, L. J. McIver, K. N. Thompson, M. Zolfo, P. Manghi, L. Dubois, K. D. Huang, A. M. Thomas, W. A. Nickols, G. Piccinno, E. Piperni, M. Punčochář, M. Valles-Colomer, A. Tett, F. Giordano, R. Davies, J. Wolf, S. E. Berry, T. D. Spector, E. A. Franzosa, E. Pasolli, F. Asnicar, C. Huttenhower, N. Segata, Extending and improving metagenomic taxonomic profiling with uncharacterized species using MetaPhlAn 4. Nat Biotechnol 41, 1633–1644 (2023).

57. H. Mallick, A. Rahnavard, L. J. McIver, S. Ma, Y. Zhang, L. H. Nguyen, T. L. Tickle, G. Weingart, B. Ren, E. H. Schwager, S. Chatterjee, K. N. Thompson, J. E. Wilkinson, A. Subramanian, Y. Lu, L. Waldron, J. N. Paulson, E. A. Franzosa, H. C. Bravo, C. Huttenhower, Multivariable association discovery in population-scale meta-omics studies. PLoS Comput Biol 17, e1009442 (2021).

58. Y. Zhou, B. Zhou, L. Pache, M. Chang, A. H. Khodabakhshi, O. Tanaseichuk, C. Benner, S. K. Chanda, Metascape provides a biologist-oriented resource for the analysis of systems-level datasets. Nat Commun 10, 1523 (2019).

59. V. Matyash, G. Liebisch, T. V. Kurzchalia, A. Shevchenko, D. Schwudke, Lipid extraction by methyl-tert-butyl ether for high-throughput lipidomics. J Lipid Res 49, 1137–1146 (2008).

60. C. G. Vasilopoulou, K. Sulek, A.-D. Brunner, N. S. Meitei, U. Schweiger-Hufnagel, S. W. Meyer, A. Barsch, M. Mann, F. Meier, Trapped ion mobility spectrometry and PASEF enable in-depth lipidomics from minimal sample amounts. Nat Commun 11, 331 (2020).

61. D. McCloskey, J. A. Gangoiti, B. O. Palsson, A. M. Feist, A pH and solvent optimized reverse-phase ion-paring-LC–MS/MS method that leverages multiple scan-types for targeted absolute quantification of intracellular metabolites. Metabolomics 11, 1338–1350 (2015).

62. G. Cao, Z. Song, Y. Hong, Z. Yang, Y. Song, Z. Chen, Z. Chen, Z. Cai, Large-scale targeted metabolomics method for metabolite profiling of human samples. Anal Chim Acta 1125, 144–151 (2020).

63. B. R. Groveman, B. Schwarz, E. Bohrnsen, S. T. Foliaki, J. A. Carroll, A. R. Wood, C. M. Bosio, C. L. Haigh, A PrP EGFR signaling axis controls neural stem cell senescence through modulating cellular energy pathways. J Biol Chem 299, 105319 (2023).

