## Supplementary Information for "Ketogenic diet synergistic reprogramming of both host and microbiome promotes tissue regeneration"

### Materials and methods

*Mice*

Conventional Specific Pathogen Free (SPF) wild-type C57BL/6, CD45.1 (Taconic line 8478), *Il1r1$^{-/-}$ (Taconic Line 8506), Il17a$^{-/-}$ (Taconic Line 8434) mice* were obtained through the NIAID-Taconic exchange program and maintained at NIAID animal facilities. *Il17ra$^{-/-}$(B6.Cg-Il17ra$^{tm2.2Koll}$/J) mice* were purchased from The Jackson Laboratories. All mice were bred and maintained under specific pathogen-free conditions at an American Association for the Accreditation of Laboratory Animal Care (AAALAC)-accredited animal facility at the NIAID and housed in accordance with the procedures outlined in the Guide for the Care and Use of Laboratory Animals. All experiments were performed at the NIAID under an animal study proposal (LHIM-3E) approved by the NIAID Animal Care and Use Committee. Mice were group housed (4–5 mice of same sex per cage) and randomly assigned to each experimental group. Unless otherwise noted, sex- and age-matched mice between 8 and 12 weeks of age were used for each experiment.

*Generation of Vg4/Vg6 double knockout (DKO) mice*

*Tcrg-V4* deficient mice were generated using CRISPR-Cas9 mediated genome editing. To generate CRISPR guide RNA (gRNA) targeting *Tcrg-V4*, the oligos 5'-TAGGTTCCCAGGGAAGATATAA-3' and 5'-AAACTTATATCTTCCCTGGGAAC-3' were annealed and ligated into the pT7-gRNA vector (Addgene). Linearized DNA was used as a template for in vitro transcription (IVT) using the MEGAshortscript T7 Transcription Kit (Thermo Fisher Scientific), which yielded the gRNAs. Cas9 mRNA was generated by using linearized pT3TS-nCas9n (Addgene) as the template for IVT with the mMESSAGE mMACHINE T3 Transcription Kit (Thermo Fisher Scientific). Both the Cas9 mRNA and the gRNAs were purified using the MEGAclear Transcription Clean-Up Kit (Thermo Fisher Scientific) and eluted in RNase-free water. C57BL/6NCr blastocysts were microinjected with Cas9 mRNA (100 ng/mL) and gRNA (100 ng/ mL). Progeny were screened by polymerase chain reaction (PCR) amplification with primers 5'-GAAGAACCCTGGCTCACAAG-3' and 5'-GGTAGGGGTTGCAAGGACAT-3', followed by Sanger sequencing with the primer 5'-AAGCCAACCTGGCAGATG-3'. The confirmed founders were bred to C57BL/6NTac mice to propagate the strain.

*Tcrg-V6*–deficient mice were generated using a similar approach. Two gRNAs targeting *Tcrg-V6* were designed using the oligonucleotides 5'-TAGGCAGTCTCACGTCACCTCT-3' and 5'-AAACAGAGGTGACGTGAGACTGC-3' (gRNA 1), and 5'-TAGGGGGTCATATGTCATCAAG-3' and 5'-AAACCTTGATGACATATGACCC-3' (gRNA 2). gRNAs and Cas9 mRNA were generated, purified, and microinjected into C57BL/6NCr blastocysts as described above. Progeny were screened by PCR with primers 5'-TGATTTCAGTGCCTCTCCTG-3' and 5'-CACAGCATTCTTTGGAGACG-3', followed by Sanger sequencing with primer 5'-TGATTTCAGTGCCTCTCCTG-3'. The confirmed founders were bred to C57BL/6NTac mice to propagate the strain.

To generate Vg4/Vg6 DKO mice, the individual knockout strains were intercrossed.

*Commensal culture and colonization*

*Staphylococcus epidermidis*1457 (*49*) and 1457 *Δsph* (*17*) were kindly provided by Dr. Michael Otto (Laboratory of Bacteriology, National Institute of Allergy and Infectious Diseases). *Staphylococcus epidermidis* NIHLM087 (*50*), NIHLM087 *ΔribD* (*21*), 1457 and 1457 *Δsph* were cultured in Tryptic Soy Broth (TSB) at 37°C until reaching OD600nm~0.8. For colonization with commensal microbes, as before (*30*), each mouse was topically associated by placing 5 ml of culture suspension (approximately $10^9$ CFU/ml) across the entire skin surface (approximately 36 $cm^2$) using a sterile swab. Application of commensal microbes was repeated every other day a total of four times. Skin tissue was analyzed 14 days after initial colonization, unless otherwise indicated. To quantify *S. epidermidis* colonization, the ear skin single cell suspension was prepared as described below (see Skin tissue processing section). Singel cell suspension was diluted in sterile PBS to perform 10 fold serial dilution. Samples were seeded on blood agar and plates and incubated at 37°C under aerobic conditions for 18 hours for CFU counts.

*Back-skin wounding and epifluorescence microscopy of back-skin wounds*

Tissue wounding and quantitation of wound healing were performed as previously described (*51*). Briefly, male mice in the telogen phase of the hair cycle were anesthetized and punch biopsies performed on back skin. Dorsal hair was shaved with clippers, and a 6-

mm biopsy punch was used to partially perforate the skin. Iris scissors were then used to cut epidermal and dermal tissue to create a full thickness wound in a circular shape. Back-skin tissue was excised 5 days after wounding, fixed in 4% paraformaldehyde in PBS, incubated overnight in 30% sucrose in PBS, embedded in OCT compound (Tissue-Tek), frozen on dry ice, and cryo-sectioned (20-μm section thickness). Sections were fixed in 4% paraformaldehyde in PBS, rinsed with PBS, permeabilized with 0.1% Triton X-100 in PBS (Sigma-Aldrich), and blocked for 1 hour in blocking buffer (2.5% Normal Goat Serum, 1% BSA, 0.3% Triton X-100 in PBS). Primary antibody to Keratin 14 (chicken, Poly9060, 1:400, Biolegend) was diluted in blocking buffer with rat gamma globulin and anti-CD16/32 and incubated overnight. After washing with PBS, a secondary antibody conjugated with Alexa647 (goat anti-chicken, Jackson ImmunoResearch) was added for 1 hour at room temperature. Slides were washed with PBS, counterstained with DAPI and mounted in Prolong Gold. Wound images were captured with a Leica DMI 6000 widefield epifluorescence microscope equipped with a Leica DFC360X monochrome camera. Tiled and stitched images of wounds were collected using a 20×/0.4NA dry objective. Images were analyzed using Imaris software (Bitplane).

*In vivo treatments*

IL-1R was blocked with a specific antagonist Kineret (Biovitrum); mice were injected intraperitoneally (*i.p.*) with Kineret or PBS control, 250 mg/kg, for 7 days starting at the time of bacterial mono-association (*30*). Imiquimod (IMQ) was topically associated on each ear pinnae. 10 mg of 5% IMQ-containing cream (Aldara Cream 5%; 3M Health Care) was applied for 5 days consecutive days. Ear thickness was measured with a digimatic caliper (Mitutoyo).

*Skin tissue processing*

Single cell suspensions from ear skin were obtained as described previously. Briefly, mice were euthanized with $CO_2$, and ear pinna skin was split into dorsal and ventral sheets and incubated in RPMI 1640 media supplemented with 2mM L-glutamine, 1mM sodium pyruvate, 1mM non-essential amino acids, 50mM b-mercaptoethanol, 20mM HEPES, 0.5mg/ml DNAse-I (Sigma-Aldrich) and 0.25mg/ml of Liberase TL (Roche) purified enzyme blend (Roche) for 1 hour and 45 min at 37°C/5% $CO_2$. Digested ears were homogenized using the Medicon/Medimachine tissue homogenizer system (Becton Dickinson) and filtered through a 70mm cell strainer.

*Flow cytometry analysis*

Fluorophore-conjugated antibodies that were used are listed in table S1. 5-OP-RU–loaded mMR1 tetramer and PBS57-loaded mCD1d tetramer were provided by the NIH Tetramer Core Facility and cells were stained in RPMI complete media for 1 hour at room temperature. Single cell suspensions were incubated for 30 minutes at 4°C with surface marker fluorophore-conjugated antibodies in 20% Brilliant Stain Buffer (BD) in PBS, in presence of purified anti-mouse CD16/32 (FcBlock, clone 93). Dead cells were excluded using a LIVE/DEAD Fixable Blue Dead Cell Stain Kit (Invitrogen). Cells were fixed and permeabilized utilizing the Foxp3/Transcription Factor Staining Buffer Set (eBioscience) and stained with fluorophore-conjugated intracellular antibodies for at least 45 minutes at room temperature.

For IL-17A detection, single cell suspensions were culture *ex vivo* at 37°C for 2.5 hours in 10% FBS supplemented (as above) RPMI 1640 media, with 5 μg/mL Ionomycin (Sigma-Aldrich), 1:1000 dilution of GolgiPlug (BD Biosciences), and 50 ng/mL phorbol myristate acetate (PMA) (Sigma-Aldrich). Cells were intracellularly stained, as described above, with anti-IL-17A BV421-conjugated antibody (Biolegend).

*SCENITH*

SCENITH was performed as described before (*31*). SCENITH reagents kits (inhibitors, puromycin, and antibodies) were kindly provided from Dr. Rafael J Argüello. For ex vivo analysis, single cell suspensions from ear skin were prepared as described above and resuspended in 10% FBS supplemented RPMI 1640 media and plated in a 96-well plate at $1.5 \times 10^6$ cells/ml. Cells were pretreated with 2-DG (50 mM), oligomycin (1 μM), or in combination for 15 min, followed by treatment with puromycin (10 μg/ml) for 45 min at 37°C. After drug treatments, samples were surface-stained and intracellularly stained with anti-puromycin (1:500; clone R4743L-E8). Glucose dependence and mitochondrial dependence were calculated as previously described (*31*).

For standard deviation calculation:
Co = MFI of anti-Puro-Fluorochrome upon Control treatment
DG = MFI of anti-Puro-Fluorochrome upon 2-Deoxy-D-Glucose treatment
O = MFI of anti-Puro-Fluorochrome upon Oligomycin A treatment
DGO = MFI of anti-Puro-Fluorochrome upon DG+O treatment

Glucose dependence (%) = (Co – DG)/(Co – DGO) x 100
Mitochondrial dependence (%) = (Co – O)/(Co – DGO) x 100

*Specific diet and dietary restriction*

Specific diets that were used are listed in table S2. All diets were purchased from Envigo Teklad Diets. Mice were fed for 4-6 weeks as described in figure legends.
Dietary restriction (DR) was performed as previously described (*3*). The daily intake of regular chow (LabDiet Advanced Protocol PicoLab Verified – 75 IF; 20% protein, 5% fat) by individually caged mice was determined by our group previously. From this, individual mice consumed approximately 2.75 g of food per day. This equates to an intake of roughly 11.4 kcal per day. As such, we provided 1.375 g of food to mice daily (5.7 kcal per day) to ensure 50% DR. Mice on DR consumed all the food provided.

*16sRNA library preparation, sequencing and analysis*

DNA was extracted from skin swab samples using a modified version of MasterPure Yeast DNA Purification kit (LCG, Bioresearch Technologies). Dual-indexing amplification and sequencing approach was taken to assess the composition of microbial communities from the given samples targeting the V1-3 hypervariable region of the 16S ribosomal RNA gene (*16S rRNA*), using primers 27F and 534R. Libraries were then quantified using the KAPA library quantification kit (Roche), pooled at equimolar concentrations and sequenced on a MiSeq system (Illumina). 16S rRNA sequencing data was analyzed with DADA2 (*52*) using the Nephele platform to generate ASVs (*53*), and the SILVA databases was utilized for taxonomic classification (*54*). We built a custom database to include the strain of interest.

*Metatranscriptome library preparation, sequencing and analysis*

Skin microbes were sampled by swabbing with two polyester swabs moistened with PBS + 10% Tween 20 and swabbed over total skin surface. RNA was isolated from skin swab heads by bead homogenization using the RNeasy Power Microbiome Kit (QIAGEN). Ribosomal RNA was depleted using the Ribocop rRNA Depletion kit (Lexogen) using META primers (Lexogen). RNA libraries were prepared using the Corall RNA-Seq V2 Library Prep Kit (Lexogen), using 20 PCR cycles, as determined by qPCR using the PCR Add on and Reamplification Kit V2 for Illumina (Lexogen). Libraries were sequenced at 750pm using 100bp paired end sequencing on Illumina NexSeq 1000 using a P2 200 cycles kit (Illumina). Metatranscriptomics data was analyzed using the Humann3 pipeline (*55*) with the metaphlan database (v.Oct22) (*56*). Outputs were then analyzed for differential expression using Maaslin 2 (*57*).

*Whole tissue RNA-seq*

2 mm punch biopsy of ear skin was submerged in RNAlater (Sigma-Aldrich) and stored at −20°C. Total tissue RNA was isolated from skin tissue using the RNeasy Fibrous Tissue Mini kit (Qiagen), as per manufacturer's instructions. Sequencing libraries were generated using the Illumina Stranded Total RNA Prep, Ligation with Ribo-Zero Plus kit, according to the manufacturer's instruction. Libraries were quantified using an Agilent Tapestation (High Sensitivity D1000 ScreenTape) and Qubit (Thermo Fisher Scientific). Libraries were sequenced as 1 x 150bp reads on an Illumina NextSeq 2000 using the P3 200 cycle kit. Sequencing reads were mapped to the C57BL/6 mouse genome (GRCm38: mm10) and differential gene expression was calculated utilizing HOMER's getDifferentialExpression with default parameters. Differentially expressed genes (FDR < 0.01, log2FC > 1) were used for gene ontology (GO) enrichment analysis with MetaScape (*58*).

*Bulk RNA-seq*

$V\gamma6^+$ $\gamma\delta$T cells were sorted in a Sony MA900 sorter from ear skin. Samples were staining with antibodies against CD45, CD90.2, TCRb, TCRgd, Vγ1, Vγ4, and Vγ5. $V\gamma6^+$ $\gamma\delta$T cells (Live (DAPI$^-$) CD45$^+$ CD90.2$^+$ TCRb$^-$ TCRgd$^{mid}$ Vγ1$^-$ Vγ4$^-$ Vγ5$^-$) were sorted directly into lysis buffer. RNA was purified using RNeasy Micro Kit (Qiagen) and libraries were prepared using Revelo High Sensitivity RNA-seq library preparation kit from Tecan according to manufacturer's protocol. Libraries were sequenced as 1 x 150bp reads on an Illumina NextSeq 2000 using the P3 200 cycle kit. Sequencing reads were mapped to the C57BL/6 mouse genome (GRCm38: mm10) and differential gene expression was calculated utilizing HOMER's getDifferentialExpression with default parameters and genes (FDR < 0.01, log2FC > 1) were considered significant. Pathway and upstream regulator analyses were conducted using Ingenuity Pathway Analysis (QIAGEN IPA). Differentially expressed genes (gene symbol, log2 fold change, adjusted p-value) were uploaded to IPA and analyzed using the Core Analysis workflow utilizing a log2FC of 1/-1 and adjusted p-value of 0.01.

*Non-targeted lipidomic analysis*

The isolation of epidermis from the underlying dermis was performed on approximately 1 $cm^2$ sections of mouse tail skin. The tissue was incubated for 30 min in 500 CU Dispase (Becton Dickinson) prepared in HBSS lacking calcium and magnesium. Following digestion, the epidermal layer was carefully peeled from the dermis using curved forceps. After a PBS wash, the epidermis was placed in methanol and stored at -80°C. Epidermal lipids were extracted as previously described (*59*). Briefly, frozen samples in 225 µl of cold methanol were mechanically homogenized with a glass beads and Precellys 24 (Bertin Technologies). A liquid-liquid extraction was then performed by adding 750 µl of cold methyl tert-butyl ether and shaking the mixture for 10 min. The addition of 188 µl of room-temperature LC-MS-grade water and a 20 seconds vertex induced phase separation. Following centrifugation at 14,000 x g for 2 min, approximately 700 µL of the resulting of upper organic layer was collected and dried under vacuum (SpeedVac, Thermo Fisher Scientific), and the resulting lipid film was reconstituted in 150 µl of 90:10 (v/v) methanol:chloroform mix for subsequent instrumental analysis. The extracted lipids were analyzed on a LC/MS system with ultra-performance liquid chromatography and trapped ion mobility spectrometry Time-of-Flight (TIMS-TOF Pro, Bruker) (*60*). Lipidomics data were analyzed by using Metaboscape 2023b (Bruker).

*Targeted metabolomic analysis*

The isolation of epidermis from the underlying dermis was performed on approximately 1 $cm^2$ sections of mouse tail skin. The tissue was incubated for 30 min in 500 CU Dispase (Becton Dickinson) prepared in HBSS lacking calcium and magnesium. Following digestion, the epidermal layer was carefully peeled from the dermis using curved forceps. After a PBS wash, the epidermis was placed in 800 µl of 50:50 (v/v) methanol:water mix and stored at -80°C. Frozen samples in 800 µl of cold methanol:wtaer were mechanically homogenized with a glass beads and Precellys 24 (Bertin Technologies). A liquid-liquid extraction was then performed by adding 400 µl of cold chloroform (Thermofisher) and shaking the mixture for 20 min at 4°C. Spin samples at max speed in a benchtop microcentrifuge (>10,000 x g) for 20 minutes at 4 4°C to induce phase separation. The resulting of upper layer was collected for subsequent instrumental analysis.

For all liquid chromatography mass spectrometry (LCMS) methods LCMS grade solvents were used.

Tributylamine was purchased from Millipore Sigma. LCMS grade water, methanol, isopropanol and acetic acid were purchased through Fisher Scientific. Aqueous metabolites were analyzed using a LD40 XR UHPLC (Shimadzu Co.) system for separation and a 6500+ QTrap mass spectrometer (AB Sciex Pte. Ltd.) for detection with polarity switching. Metabolites were separated on a Water XBridge Amide column (3.5 µm, 3 mm X 100 mm) and eluted using a binary gradient from 100% 5mM ammonium acetate acid in 5% water pH adjusted to 7.5 and 95% acetonitrile to 100% 5 mM ammonium acetate 95% water pH adjusted to 7.5 and 5% acetonitrile over 6.75 minutes. Multiple reaction monitoring ion pairs (MRM) for metabolite detection were adopted from previous publications with multiple MRMs used for signals with convoluting peaks (*61–63*).

All signals were integrated using SciexOS 3.1 (AB Sciex Pte. Ltd.). Signals were inspected visually for presence and signals with greater than 50% missing values were discarded. Any remaining missing values were replaced with the minimum detected value within the dataset. Metabolites with multiple MRMs were quantified with the higher signal to noise MRM. Filtered datasets were total sum normalized after initial filtering. A Benjamini-Hochberg method for correction for multiple comparisons was imposed where indicated.

*Statistical analysis*

Statistical methods utilized are described in figure legends. Groups were compared using Prism software (version 10).

**fig. S1**

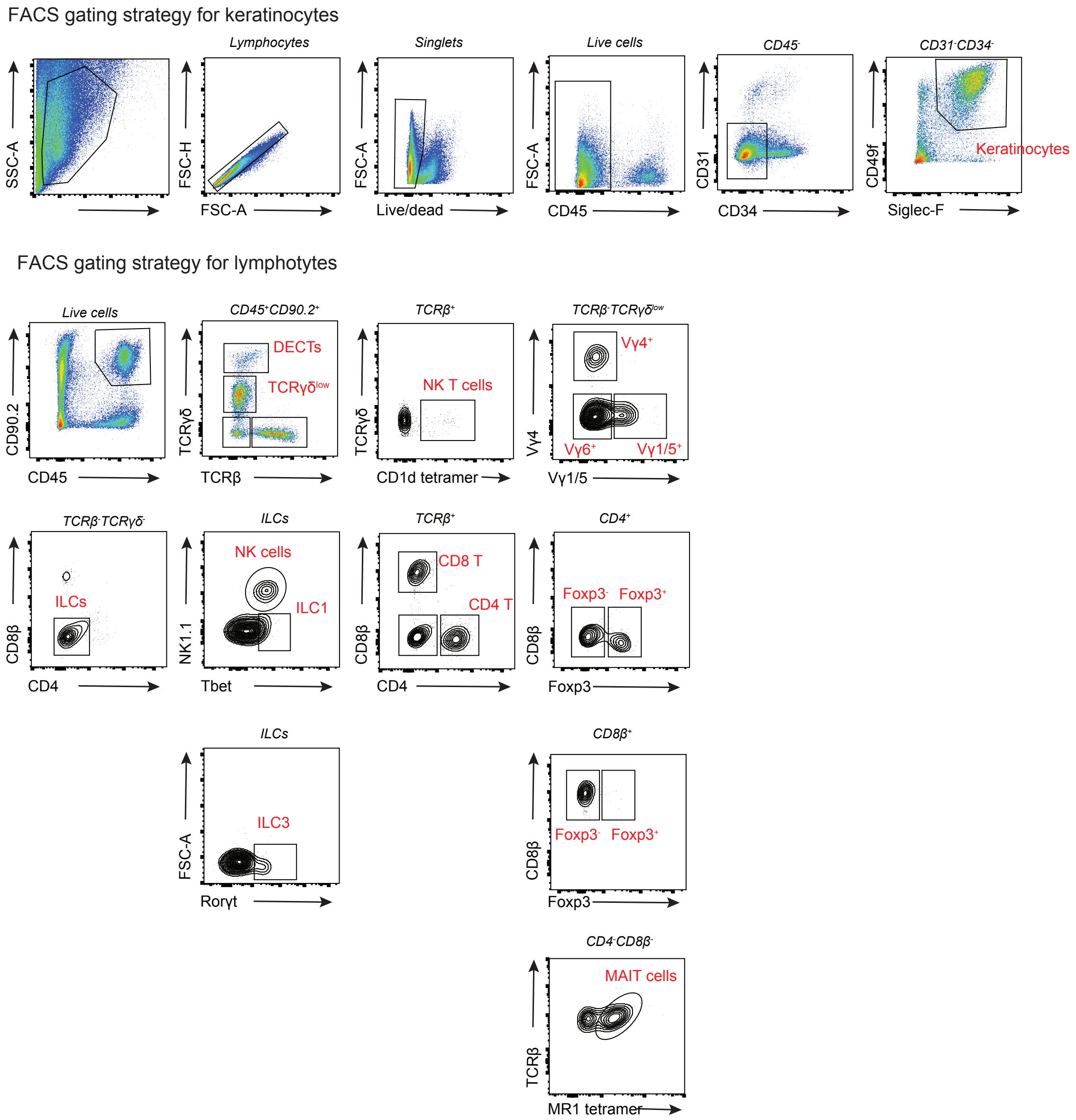


**fig. S1. Flow cytometry gating strategies.**

Representative flow cytometry plots showing the gating strategy used to define keratinocytes and lymphocyte populations from ear skin.

**fig. S2**

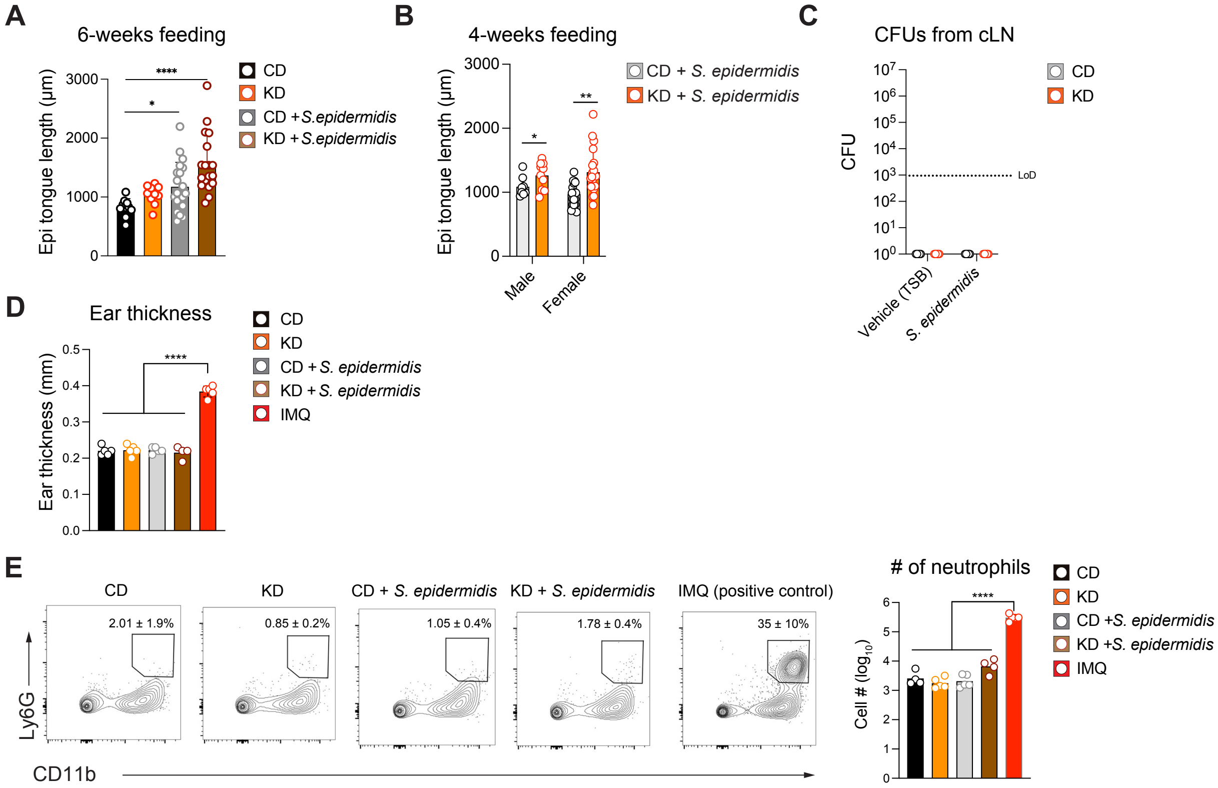


**fig. S2. KD + *S. epidermidis* does not induce skin inflammation.**

(A–E) WT mice were fed a CD or KD for a total of six weeks (A) or four weeks (B–D) and were topically associated with *S. epidermidis* or TSB.

(A, B) Epidermal tongue length was measured 5 days after punch biopsy.

(C) Colony-forming units (CFU) of *S. epidermidis* recovered from skin–draining cervical lymph nodes.

(D, E) Mice were topically treated with imiquimod (IMQ) for five days as a positive control for skin inflammation. (D) Ear thickness measured by caliper. (E) Representative flow cytometry plots (left) showing frequencies of live $CD45^+$ $CD11b^+$ $Ly6G^+$ neutrophils, and bar graphs (right) showing absolute neutrophil numbers per ear pinna.

Panels (A, B) show pooled data from at least three independent experiments; (C–E) are representative of two independent experiments. Data are mean ± s.e.m.; each dot represents an individual epidermal tongue measurement (A, B) or mouse (C–E). (A) One-way ANOVA followed by Dunnett's test; (B) one-way ANOVA followed by multiple t-tests;

(D, E) one-way ANOVA followed by Tukey’s test. $^{*}P < 0.05$; $^{**}P < 0.01$; $^{***}P < 0.001$; $^{****}P < 0.0001$

**fig. S3**

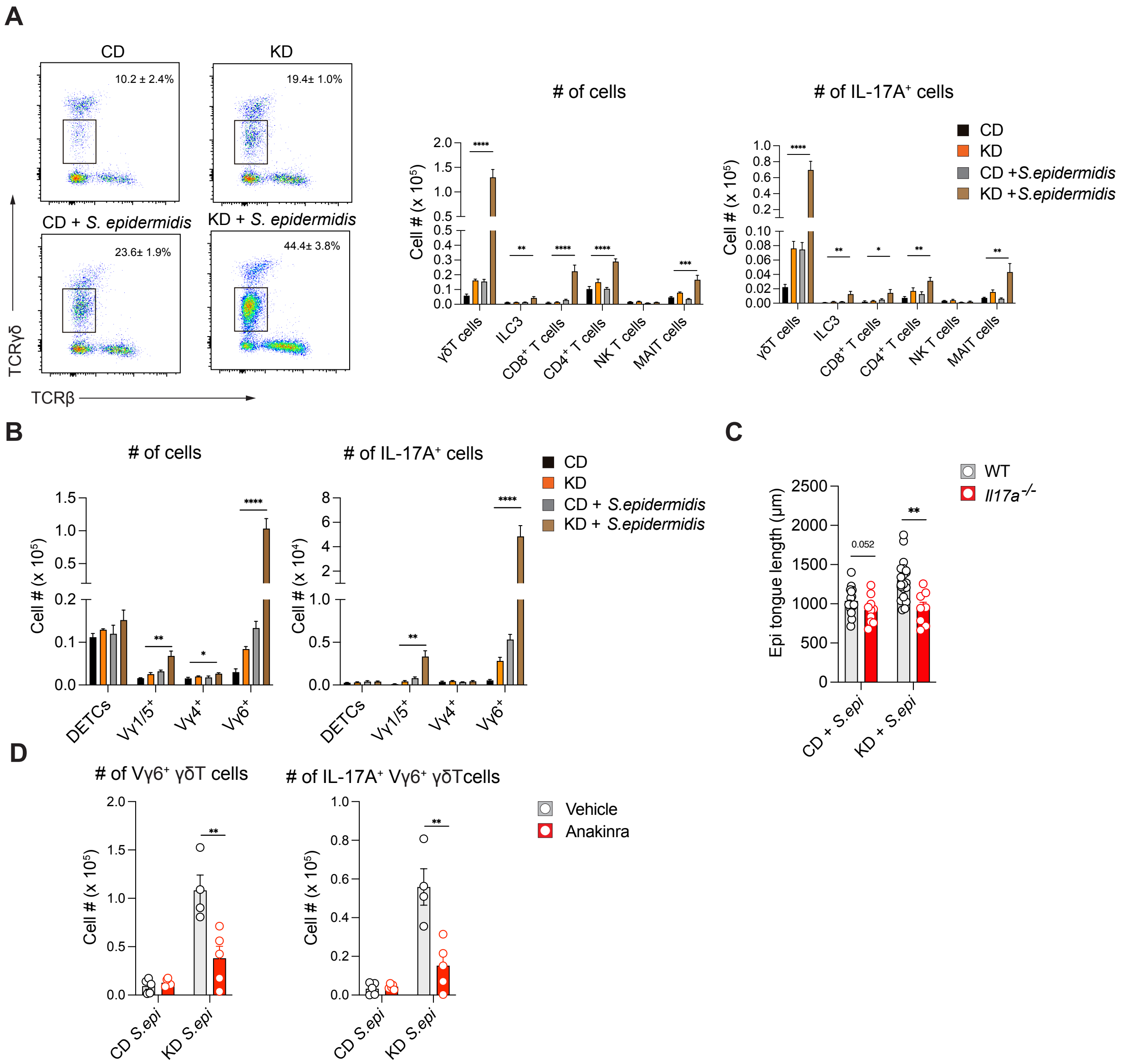


**fig. S3. KD + *S. epidermidis* enhances wound healing through activation of skin γδ T cells.**

(A–D) WT (A, B, D), *Il17a*$^{-/-}$, or age-matched control mice (C) were fed a CD or KD for four weeks and were topically associated with *S. epidermidis* or TSB.

(A) Representative flow cytometry plots (left) showing frequencies of γδ T cells (gated on live CD45$^{+}$ CD90.2$^{+}$ TCRβ$^{-}$ γδTCR$^{low}$) in ear pinna, and bar graphs (right) showing absolute γδ T cell numbers.

(B) Absolute numbers of indicated cells in ear pinna.

(C) *Il17a*$^{-/-}$ and WT mice were wounded on the back skin, and epidermal tongue length was measured 5 days post-injury.
(D) WT mice were treated with anakinra or vehicle for 7 days to block IL-1 signaling; bar graphs show absolute numbers of indicated cells in ear pinna.

Panels (A, B, D) show representative data from at least two independent experiments; panel (C) shows pooled data from two independent experiments. Data are mean ± s.e.m.; each dot represents an individual mouse or epidermal tongue length. (A, B) One-way ANOVA followed by Dunnett's test; (C, D) one-way ANOVA followed by multiple t-tests. *p < 0.05; **p < 0.01; ***p < 0.001; ****p < 0.0001.

**fig. S4**

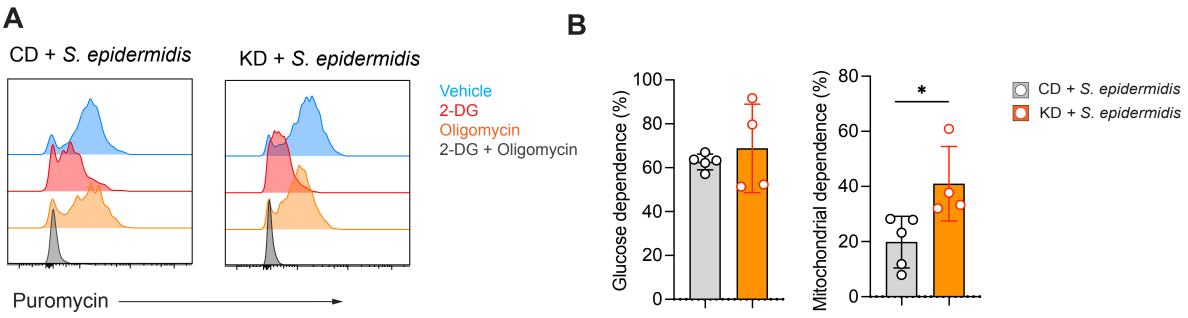


**fig. S4. KD feeding alters metabolic states of skin γδ T cells.**

WT mice were fed a CD or KD for four weeks and topically associated with *S. epidermidis*. Single-cell suspensions were prepared from ear pinna, and cellular metabolic states were assessed using the SCENITH kit.

(A) Representative histograms showing puromycin incorporation in Vγ6+ γδ T cells following treatment with vehicle, 2-deoxyglucose (2-DG; glycolysis inhibitor), oligomycin (OXPHOS inhibitor), or 2-DG + oligomycin, and subsequent incubation with puromycin for 45 min.

(B) Bar graphs showing calculated glucose dependence and mitochondrial dependence in Vγ6+ γδ T cells.

Data are shown as mean ± s.e.m.; each dot represents an individual mouse. Statistical significance was determined using Student's t-test. *$P < 0.05$.

**fig. S5**

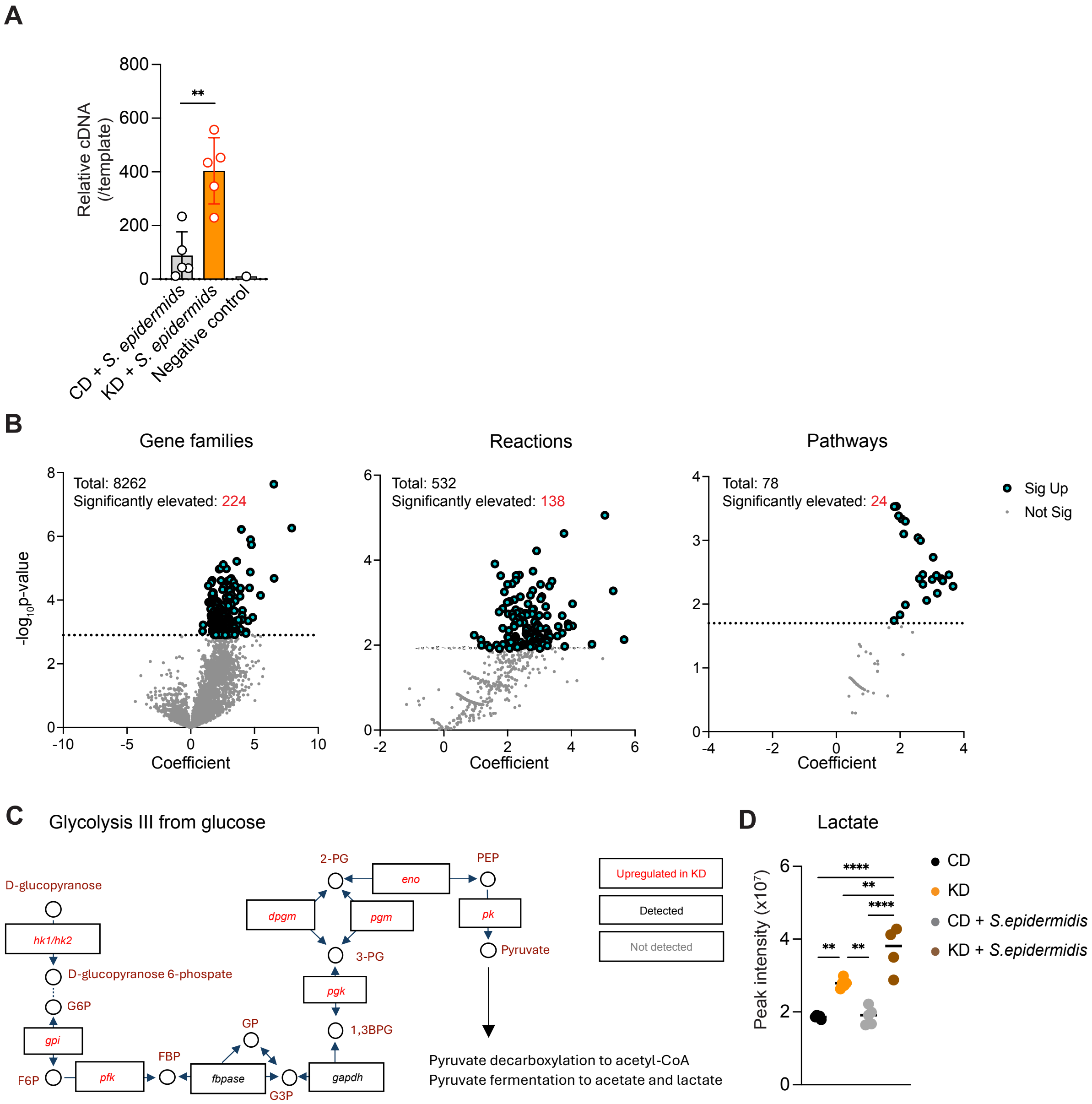


**fig. S5. Skin microbiota adapt to KD-induced environmental changes.**

(A–D) WT mice were fed a CD or KD for four weeks and were topically associated with *S. epidermidis* or TSB.

(A–C) Metatranscriptomic analysis of back-skin swabs. Bacterial RNA was extracted, rRNA was depleted, and cDNA was generated by random-primed reverse transcription. cDNA input was quantified by qPCR and normalized before library preparation.

(A) Amount of bacterial cDNA obtained from each swab; brand-new swabs served as negative controls.
(B) Volcano plots showing differentially expressed gene families (left), annotated reactions (middle), and enriched pathways (right) between CD + *S. epidermidis* and KD + *S. epidermidis*. A total of 8,262 gene families, 532 reactions, and 76 pathways were detected, of which 224 gene families, 138 reactions, and 24 pathways were significantly upregulated in the KD group. Features enriched in KD + *S. epidermidis* are shown in blue.
(C) Differential expression of glycolysis-related bacterial genes; genes significantly upregulated in KD + *S. epidermidis* are shown in red.
(D) Epidermal concentration of lactate measured by targeted metabolomics.

Abbreviations and EC numbers: *hk1/hk2, hexokinase 1/2* (EC 2.7.1.1/EC 2.7.1.2); *gpi, glucose-6-phosphate isomerase* (EC 5.3.1.9); *pfk, 6-phosphofructokinase* (EC 2.7.1.11); *fbpase, fructose-bisphosphate aldolase* (EC 4.1.2.13); *gapdh, glyceraldehyde-3-phosphate dehydrogenase* (EC 1.2.1.12); *pgk, phosphoglycerate kinase* (EC 2.7.2.3): *dpgm, 2,3-diphosphoglycerate-dependent phosphoglycerate mutase* (EC 5.4.2.11), *pgm, 2,3-diphosphoglycerate-independent phosphoglycerate mutase* (EC 5.4.2.12); *eno, enolase* (EC 4.2.1.11); *pk, pyruvate kinase* (EC 2.7.1.40).
Metabolite abbreviations: G6P, glucose-6-phosphate; F6P, fructose-6-phosphate; FBP, fructose-1,6-bisphosphate; GP, glycerate 3-phosphate; G3P, glyceraldehyde 3-phosphate; 1,3-BPG, 1,3-bisphosphoglycerate; 3-PG, 3-phosphoglycerate; 2-PG, 2-phosphoglycerate; PEP, phosphoenolpyruvate.

Each dot represents one mouse. (A) Student's t-test; (D) one-way ANOVA followed by Tukey's test. *$P < 0.05$; **$P < 0.01$; ****$P < 0.0001$.

**fig. S6**

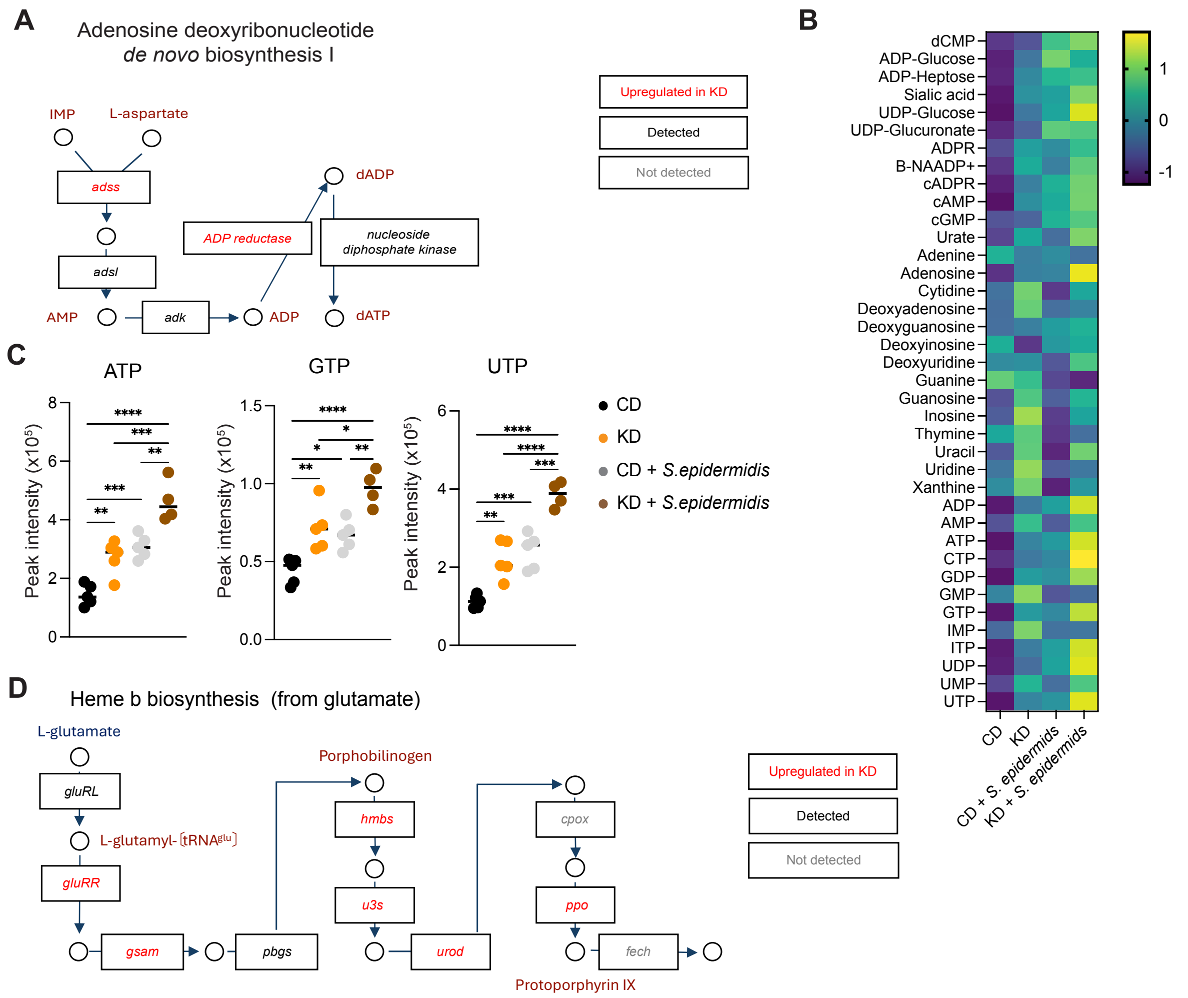


**fig. S6. KD feeding enhances nucleotide and porphyrin production by *S. epidermidis*.**

WT mice were fed a CD or KD for four weeks and topically associated with *S. epidermidis* or TSB.

(A, D) Changes in gene expression related to adenosine deoxyribonucleotide *de novo* biosynthesis (A) and heme b biosynthesis (D) in skin bacteria from CD + *S. epidermidis* and KD + *S. epidermidis* mice. Genes significantly upregulated in the KD + *S. epidermidis* group are shown in red. Data were obtained from skin metatranscriptomics (related to Fig. 4A and fig. S5).

(B, C) Nucleotide concentrations were measured by targeted metabolomics. (B) Heat map showing Z-scored nucleotide concentrations in the epidermis. (C) Dot plots showing nucleotide peak intensities.

Abbreviations and EC numbers: *adss, adenylosuccinate synthase* (EC 6.3.4.4); *adsl, adenylosuccinate* lyase (EC 4.3.2.2); *adk, adenylate kinase* (EC 2.7.4.3); *ADP reductase, adenosine diphosphate reductase* (EC 1.17.4.1*); nucleoside diphosphate kinase* (EC 2.7.4.6); *gluRL, glutamate–tRNA ligase* (EC 6.1.1.17*); gluRR, glutamyl tRNA reductase* (EC 1.2.1.70); *gsam, glutamate-1-semialdehyde 2,1-aminomutase* (EC 5.4.3.8); *pbgs, porphobilinogen synthase* (EC 4.2.1.24); *hmbs, hydroxymethylbilane synthase* (EC 2.5.1.61); *u3s, uroporphyrinogen III synthase* (EC 4.2.1.75*); urod, uroporphyrinogen decarboxylase* (EC 4.1.1.37); *cpox, coproporphyrinogen oxidase* (EC 1.3.3.3); *ppo, protoporphyrinogen oxidase* (EC 1.3.3.4); *fech, protoporphyrin ferrochelatase* (EC 4.98.1.1). Metabolite abbreviations: IMP, inosine monophosphate; AMP, adenosine monophosphate; ADP, adenosine diphosphate; dADP, deoxyadenosine diphosphate; ATP, adenosine triphosphate; dATP, deoxyadenosine triphosphate; GTP, guanosine triphosphate; UTP, uridine triphosphate.

(C) Each dot represents one mouse; one-way ANOVA followed by Tukey's test. *P < 0.05; **P < 0.01; ****P < 0.0001

**fig. S7**

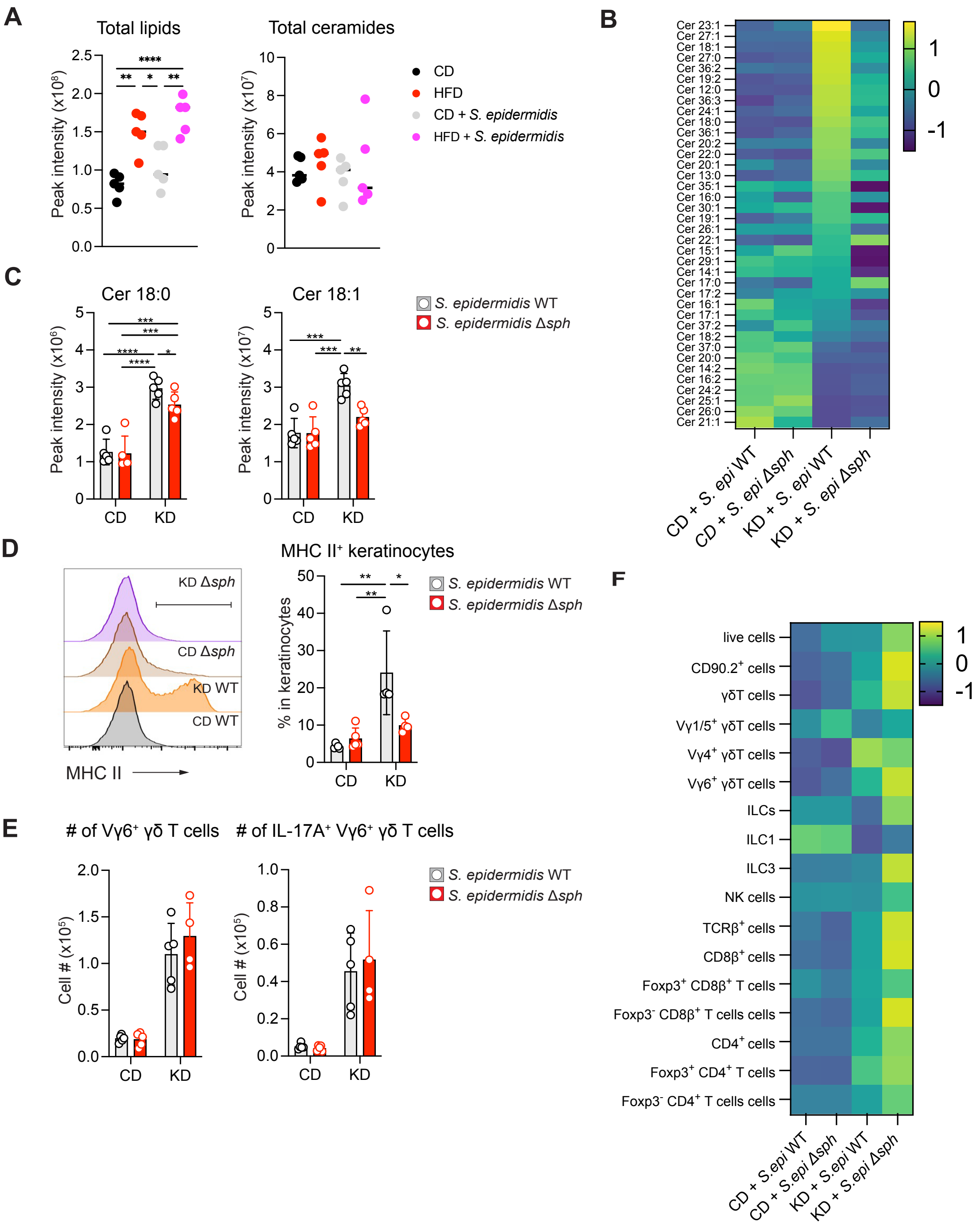


**fig. S7. Bacterial ceramides may directly activate keratinocytes.**

(A) WT mice were fed a CD or high-fat diet (HFD) for four weeks and topically associated with *S. epidermidis* or TSB. Dot plots show peak intensities of indicated lipid species in the epidermis.

(B–F) WT mice were fed a CD or KD for four weeks and topically associated with *S. epidermidis* 1457 (WT) or *S. epidermidis Δsph*.

(B, C) Ceramide concentrations were measured by lipidomics in the epidermis. (B) Heat map showing Z-scored concentrations. (C) Bar graphs showing peak intensities.

(D) Left, representative histogram of MHC II expression on keratinocytes; right, frequencies of MHC II$^+$ keratinocytes.

(E) Absolute numbers of indicated cells in ear pinna.

(F) Heat map showing Z-scored absolute numbers of lymphocyte subsets in ear pinna.

Panels (D–F) show representative data of three independent experiments. Data are mean ± s.e.m.; each dot represents one mouse. (E) One-way ANOVA followed by multiple t-tests; (A, C, D) one-way ANOVA followed by Tukey's test. *$P < 0.05$; **$P < 0.01$; ***$P < 0.001$; ****$P < 0.0001$

**table S1. Antibodies for flow cytometry**

| Antibodies | Clone | Source | Cat# |
|---|---|---|---|
| anti-mouse CD4 BV510 | RAM4/5 | Biolegend | 100553 |
| anti-mouse CD8β FITC | H35-17.2 | Invitrogen | 11-0083-82 |
| anti-mouse CD8β BUV395 | H35-17.2 | BD | 740278 |
| anti-mouse CD11b BUV737 | M1/70 | BD | 612800 |
| anti-mouse CD31 PerCP-eF710 | 390 | Invitrogen | 46-0311-82 |
| anti-mouse CD34 eF660 | RAM34 | eBioscience | 50-0341-82 |
| anti-mouse CD45 APC-eF780 | 30-F11 | Invitrogen | 47-0451-82 |
| anti-mouse CD45 BV510 | 30-F11 | Biolegend | 103138 |
| anti-mouse CD49f PE | eBioGoH3 | eBioscience | 12-0495-82 |
| anti-mouse CD90.2 BV785 | 30-H12 | Biolegend | 105331 |
| anti-mouse CD90.2 AF700 | 30-H12 | Biolegend | 105320 |
| anti-mouse NK1.1 PerCP-Cy5.5 | PK136 | Invitrogen | 45-5941-82 |
| anti-mouse Foxp3 FITC | FJK-16s | Invitrogen | 11-5773-82 |
| anti-mouse IL-17A BV421 | TC11-18H10.1 | Biolegend | 506926 |
| anti-mouse Ki67 PE-Cy7 | SolA15 | Invitrogen | 25-5698-82 |
| anti-mouse Ly6G AF700 | 1A8 | BD | 561236 |
| anti-mouse MHC II BV650 | M5/114.15.2 | Biolegend | 107641 |
| anti-mouse RORγt PE-CF594 | Q31-378 | BD | 562684 |
| anti-mouse Sca-1 FITC | D7 | Biolegend | 108106 |
| anti-mouse T-bet PE-Cy7 | 4B10 | Biolegend | 644824 |
| anti-mouse TCRβ FITC | H57-597 | eBioscience | 11-5961-85 |
| anti-mouse TCRβ PE-CF594 | H57-597 | BD | 562841 |
| anti-mouse TCRβ BUV737 | H57-597 | BD | 612821 |
| anti-mouse TCRγδ PE-CF594 | GL3 | BD | 563532 |
| anti-mouse TCRγδ BV650 | GL3 | BD | 563993 |
| anti-mouse TCR Vγ1 (Tonegawa) APC | 2.11 | Biolegend | 141108 |
| anti-mouse TCR Vγ5 (Tonegawa) APC | 536 | Biolegend | 137506 |
| anti-mouse TCR Vγ4 (Tonegawa) PerCP-Cy5.5 | UC3-10Ab | Invitrogen | 46-5828-82 |
| anti-mouse CD16/32 | 2.4G2 | BioXCell | BE0307 |
| anti-puromycin PE-CF594 | R4743L-E8 | From Dr. Rafael J Argüello Lab | N/A |

**table S2. Specific diets**

| Diet | Source | Cat# | Comment |
|---|---|---|---|
| Ketogenic diet | Inotiv | TD.160153.PWD | % kcal from protein: 9.5%, carbohydrate: 0.0%, Fat: 90.5% |
| Control diet for TD.160153.PWD | Inotiv | TD.210814 | % kcal from protein: 9.7%, carbohydrate: 77.8%, Fat: 12.5% |
| High fat diet (HFD) | Inotiv | TD.06414 | % kcal from protein: 18.3%, carbohydrate: 21.4%, Fat: 60.3% |
| Control diet for TD. 06414 | Inotiv | TD.150064 | % kcal from protein 18.6%, carbohydrate 69.1%, Fat: 10.5% |
| High cholesterol diet | Inotiv | TD.07841 | Adds 2% cholesterol to TD.00217 |
| Control diet for TD.07841 | Inotiv | TD.00217 | % kcal from protein 22%, carbohydrate 66%, Fat: 12% |
| High fiber diet | Inotiv | TD.210347 | Modified from TD.130654 with 100 g corn starch replaced by inulin. |
| Control diet for TD.210347 | Inotiv | TD.130654 | Modified AIN-93G<br>% kcal from protein 18.8%, carbohydrate 63.9%, Fat: 17.2% |
| Amino acids diet | Inotiv | TD. 210529 | Modification of TD.210528 replacing casein with the amino acids mix patterned off casein. |
| Control diet for TD.210529 | Inotiv | TD.210528 | Modified AIN-93G<br>% kcal from protein 19.0%, carbohydrate 63.6%, Fat: 17.4% |
| Vegan diet | Inotiv | TD.220273 | Modification of TD.150345 replacing casein with soy protein and increasing fiber to 29.9 g/1000 kcal with a 50:50 mix of cellulose and inulin. |
| Control diet for TD.220273 | Inotiv | TD.150345 | % kcal from protein 10%, carbohydrate 77%, Fat: 13% |
